# Comparative single-cell transcriptomic roadmap of mammalian fetal ovarian development

**DOI:** 10.64898/2026.05.01.721341

**Authors:** Yifei Fang, Shiyao Han, Junmei Zhang, Shengcan Xie, Yue Su, IISAGE Consortium, Yulei Wei, Young Tang, Jingyue Ellie Duan

## Abstract

Early ovarian development establishes the cellular basis of female reproduction, yet its cellular composition and developmental dynamics remain poorly characterized across mammalian species. Here, we generated a cross-species single-cell transcriptomic atlas of early female gonadal development in cattle (E38-E112), human (PCW6-16), and mouse (E11.5-E18.5), integrating 107,930 cells across comparable developmental windows, spanning sex determination and early ovarian differentiation. We identified 11 shared gonadal cell types across three species, including germ cells and granulosa cells, and uncovered a previously undescribed bovine-specific cell population with steroidogenic features, suggesting species differences in ovarian somatic lineage development. Epigenetic regulators exhibited dynamic activation in germ cells and granulosa cells. Developmental trajectory analysis revealed shared and species-specific genes associated with germ cell and granulosa cell development, including conserved dynamically expressed genes such as *TFAP2C* and *ZCWPW1* in germ cells and *FOS* and *JUNB* in granulosa cells. Cross-species cell type classification using a machine learning support vector machine (SVM) model revealed transcriptionally conserved cell types, such as immune cells and germ cells, while granulosa cells showed substantial cross-species divergence, associated with extracellular matrix-related gene expression. Together, our study provides a comparative framework for understanding conserved and divergent mechanisms of early ovarian development across mammals.

## Introduction

Early mammalian embryonic gonads consist of primordial germ cells (PGCs) and surrounding gonadal somatic cells. PGCs are specified in the epiblast near the proximal allantois. In humans, they begin migrating at approximately 4 weeks of development^1^ (embryonic day E7.5 in mice^2^, E20 in bovine^3^) and travel through the hindgut endoderm before colonizing the genital ridge. At this stage, the genital ridge is bipotential and has the capacity to develop into either testes or ovaries^4^. Gonadal somatic cells arise from the genital ridge at around 32 days of pregnancy in humans (E10.5 in mice, E27-E31 in bovine), in which coelomic epithelial cells differentiate into two major somatic precursor lineages, including supporting cell precursors and steroidogenic cell precursors^5^.

In males, upregulation of the *SRY-SOX9-FGF9* axis initiates male sex determination and drives somatic cells to differentiate into pre-Sertoli cells^6^. In females, in the absence of *SRY*, the RSPO1/WNT4/β-catenin and *FOXL2* pathways are activated to promote gonadal somatic cell differentiation into pre-granulosa cells^7^. As pre-granulosa cells differentiate, PGCs initiate meiosis and eventually arrest at the diplotene stage of prophase I before puberty^8^. In parallel, PGCs undergo extensive epigenetic reprogramming, characterized by global DNA demethylation and chromatin remodeling that establishes a permissive transcriptional landscape^9^.

PGC proliferation and early germ cell differentiation in females are controlled by conserved regulators such as *DAZL*^10,11^ and *DDX4*^12,13^, which are essential for germ cell maintenance and the acquisition of mitotic competence across mammals. While core features of gonadal development are broadly conserved across mammals, important species-specific regulatory mechanisms have also been reported. Meiosis is subsequently initiated by STRA8, a factor essential for meiotic entry in mice^14^, whereas in humans, meiotic initiation is associated with STRA8 expression but appears to involve a more complex regulatory network, with partial dependence on retinoic acid^15^. Despite these differences, core meiotic regulators such as *SYCP3*^13,14^ and *REC8*^16,17^ are conserved across mammals.

Early gonadal development has important implications for both human health and livestock production. Impairments in these processes can result in disorders of sex development (DSDs) or infertility in mammals^18^. For example, mutations in SRY can cause 46,XY gonadal dysgenesis and sex reversal in humans^19^. In livestock species, abnormalities in gonadal development, such as ovarian dysgenesis or ovarian hypoplasia^20^, represent the most common cause of reproductive failure. In livestock production, early gonadal development has long attracted research interest due to its direct impact on puberty onset and its potential to improve reproductive efficiency. However, limited mechanistic understanding has restricted its practical application.

Recent advances in single-cell RNA sequencing (scRNA-seq) have enabled high-resolution characterization of gonadal development in humans^21^, mice^22^, and livestock^23^, revealing both conserved and species-specific transcriptional dynamics in germ cells and somatic lineages during sex determination and ovarian differentiation. By resolving cellular heterogeneity, scRNA-seq facilitates the identification of rare or transient cell populations at a limited developmental window, such as supporting cell intermediates^21^, and the reconstruction of lineage-specific developmental trajectories for both germ cells and somatic cells. Despite these advances, cross-species comparison of gonadal development remains technically and experimentally challenging. For example, each species has differences in developmental timing, which makes it hard to align collected samples for equivalent stages. A limited number of one-to-one orthologous genes map across multiple species, which further challenges the number of genes that can be compared. Moreover, technical variability in single-cell data generation and processing from different studies further hinders integrative analyses. As a result, despite scRNA-seq having been generated in developing gonads of multiple mammalian species, a robust and systematic framework for delineating conserved and divergent regulatory mechanisms of gonadal development across mammals is still lacking.

In this study, we generated a bovine RNA-seq dataset covering six time points during early ovarian development (E38–E112) and performed a cross-species comparative scRNA-seq analysis with stage-matched human (PCW6–16) and mouse (E11.5–E18.5) datasets. Our objectives were: 1) compare the dynamics of germ and somatic cell populations across species during major events in early development, 2) identify conserved and species-specific regulators involved in germ cells and granulosa differentiation, and 3) characterize cell-cell communication between germ cells and granulosa cells during fetal ovarian development. Together, this analysis offers a comparative framework to better understand how the early female gonad is regulated at the molecular level during development across mammals, revealing conserved and species-specific regulatory programs and cellular dynamics.

## Methods

### Experimental Model and Data Availability

Human (PCW6, PCW7, PCW10, PCW12, PCW13, and PCW16) and mouse (E11.5, E12.5, E14.5, E16.5, and E18.5) single-cell RNA-seq datasets used in this study are publicly accessible from ArrayExpress (E-MTAB-10551) and GEO (GSE136441). Single-cell RNA sequencing data for fetal bovine gonads are available in the China National Center for Bioinformation (CNCB) database under accession number: OMIX012725.

### Ethics Statement

All animal procedures were conducted in accordance with the National Research Council’s Guide for the Care and Use of Laboratory Animals and were approved by the Institutional Animal Care and Use Committee (IACUC) of Northwest A&F University (protocol IACUC2024-1204, approved on 10 December 2024). Fetal tissues were obtained from a local slaughterhouse as previously described^24^.

### Fetal Bovine Gonad Collection

Estrus in donor cows was monitored daily to determine the timing of pregnancy. Artificial insemination was performed once at the onset of estrus. Uteri were collected from cows maintained under natural conditions and transported to a sterile laboratory for processing. Embryos were dissected, and fetal gonads were isolated under sterile conditions. Morphologically intact fetal gonads with sufficient cell viability were collected from female embryos at developmental stages E38, E46, E73, E83, E92, and E112.

### Single-Cell Suspension Preparation of Fetal Gonads

To isolate germ cells and gonadal somatic cells, fetal gonads were dissected under a stereomicroscope and placed into 1.5-mL tubes containing an enzymatic digestion solution (1 mg/mL Collagenase IV, 1 mg/mL Hyaluronidase, 0.25% Trypsin, and 1 mg/mL DNase I). Samples were incubated at 37°C for 30 minutes, after which serum was added to stop the enzymatic reaction. The resulting cell mixture was gently pipetted, passed through a 40-μm strainer, and resuspended in 0.1% serum/PBS to generate single-cell suspensions for 10x Genomics scRNA-seq.

### Single-Cell RNA Sequencing

Single-cell suspensions with >80% viability were adjusted to a final concentration of 700–1200 cells/μL. For each sample, single-cell encapsulation and library preparation were performed at Novogene Bioinformatics Technology Co., Ltd. (Tianjin, China) using the MobiCube® High Throughput Single Cell 3’ RNA-Seq Kit (v2.1), following the manufacturer’s instructions. Cells or nuclei were loaded into the microfluidic chip of the Chip A Single Cell Kit (MobiDrop, cat. no. S050100201) and processed on the MobiNova-100 system (MobiDrop, cat. no. A1A40001) to generate droplets. Each droplet contained a single cell or nucleus together with a gel bead carrying millions of oligonucleotides with unique cell barcodes. After droplet formation, light-mediated cleavage using the MobiNovaSP-100 (MobiDrop, cat. no. A2A40001) released the oligos into the reaction mix. mRNAs were captured by oligo(dT) within the gel beads, followed by reverse transcription and cDNA amplification. Libraries were constructed using the High Throughput Single Cell 3’ RNA-Seq Kit v2.0 (MobiDrop (Zhejiang) Co., Ltd., cat. no. S050200201) and the 3’ Single Index Kit (MobiDrop (Zhejiang) Co., Ltd., cat. no. S050300201). Sequencing was performed on an Illumina NovaSeq platform using 150-bp paired-end reads at Novogene (Tianjin, China).

### Single-Cell RNA Sequencing Data Processing

Raw sequencing data were processed using Cell Ranger (v3.1, 10x Genomics)^25^. For bovine samples generated in this study, sequencing reads were aligned to the bovine reference genome (ARS-UCD2.0) using Cell Ranger according to the manufacturer’s instructions. Human (GRCh38) and mouse (GRCm38) reference genomes were used for processing the corresponding datasets. Species-specific reference genomes were built using the Cell Ranger mkref pipeline. The filtered feature–barcode matrices generated by Cell Ranger were used for downstream analyses.

### Quality Control and Cell Type Annotation

The quality control was performed using the Seurat package (v5.2.1)^26^. Cells expressing fewer than 200 genes and genes detected in fewer than 3 cells were excluded. Cells with nFeature_RNA or nCount_RNA values outside ±3 median absolute deviations (MADs) from the median were classified as outliers and removed. Additionally, cells with mitochondrial gene expression exceeding 10% were excluded. Doublets were identified and removed using scDblFinder (v1.61.0)^27^. One-to-one orthologous genes from Ensembl were used to integrate single-cell RNA-seq datasets from bovine, human, and mouse. Data were normalized using Seurat, and the top 5,000 highly variable genes were selected for downstream analyses. During data normalization, sequencing batches, cell cycle scores, nFeature_RNA, and the percentage of mitochondrial and ribosomal genes were regressed out. For identifying cell clusters, the optimal resolution was selected with the aid of the clustree (v0.5.1) package^28^ and the dimensionality reduction was performed using the top 25 principal components. To annotate cell clusters, the FindAllMarkers function was used with default parameters to identify cluster-specific marker genes. Each cluster was manually annotated using canonical markers to identify mammalian ovarian cell types.

### Pseudotime Trajectory Analysis

To reconstruct developmental trajectories of germ cells and granulosa cells, pseudotime analysis was performed using Monocle3 (v1.3.7)^29^ following the standard workflow. Analyses were conducted separately for each species. For each cell type, cells enriched for early-stage markers were selected to define the root of the trajectory, and pseudotime values were inferred using Monocle3. Trajectory-associated genes were identified by spatial differential expression analysis using the graph_test () function (q value < 0.05). For each cell type, trajectory-associated genes were grouped into 3-5 modules along pseudotime by unsupervised hierarchical clustering and visualized using ClusterGVis (v0.1.4)^30^. Gene sets identified from each species were then compared using Venn diagrams to determine shared and species-specific trajectory-associated genes.

### Gene Ontology (GO) enrichment analysis

Trajectory-associated genes identified from the pseudotime analysis were subjected to Gene Ontology (GO) enrichment analysis using ShinyGO (v0.85.1)^31^. GO analysis was performed separately for each species.

### Transcriptional Regulatory Network Analysis

Gene regulatory network analysis was performed using pySCENIC (v0.12.1)^32^ for germ cells and granulosa cells, with analyses conducted separately for each species. Raw count matrices were used to infer transcription factor–centered gene co-expression networks using GRNBoost2. These co-expression modules were subsequently filtered by cis-regulatory motif enrichment analysis to define regulons comprising transcription factors and their predicted target genes. Regulon activity was quantified at the single-cell level using AUCell, generating regulon activity (AUC) scores. Transcription factors exhibiting conserved or species-specific regulon activity were identified during germ cell and granulosa cell developmental progression.

### Cell-Cell Communication Analysis

Cell-cell communication analysis was performed using the CellChat package (v2.1.0)^33^. Cell–cell interactions mediated by ligand–receptor pairs were inferred with the default parameters. The normalized counts were used as input.

### SVM-Based Cross-Species Cell Type Comparison

To compare cell-type transcriptomic similarity across species, a linear support vector machine (SVM)^34^ was trained on the human dataset and applied to mouse and bovine cells. To reduce class imbalance, human cell types were downsampled to match the smallest cell-type population, 305 cells. 75% of the data were used for training the model and 25% of the data were used to test the model. This procedure was repeated using bootstrapping to ensure the robustness of model performance. Raw UMI counts were normalized to counts per million (CPM) using edgeR^35^ and were log1p-transformed. The top 2,000 highly variable genes were selected based on variance in the human dataset and used as features for model training and projection. The SVM was trained using LiblineaR^36^ with annotated human cell types as labels and subsequently used to project mouse and bovine cells. For each cell, the predicted probability corresponding to its annotated cell type was extracted and summarized by cell type to assess relative cross-species similarity.

To further examine species-specific transcriptional features within granulosa cells, an additional multiclass SVM was trained on granulosa cells from human, mouse, and bovine. Species-associated genes were identified by ranking the absolute values of the learned linear coefficients for each class. In addition, to infer the identity of the unclassified bovine-specific cell population, an SVM was trained on bovine cells excluding this population and used to predict the most similar bovine cell types.

## Results

### Integration of single-cell RNA-seq data from early female gonads across bovine, human, and mouse

To characterize the cellular landscape of early female gonadal development in cattle, we generated single-cell RNA sequencing (scRNA-seq) profiles from fetal bovine ovaries across six gestational stages (**Figure 1A**). These stages encompass key developmental milestones: sex determination (E38, E46), pre-granulosa cell differentiation (E73, E83, E92, and E112), and development of early germ cells, including primordial germ cells (PGCs), meiotic germ cells and early oocytes. Here, we collectively refer to germ cells across all developmental stages, including PGCs, meiotic germ cells, and early oocytes as “germ cells”. To enable cross-species comparison, we integrated publicly available scRNA-seq datasets from human fetal ovaries (post-conception weeks PCW6, PCW7, PCW10, PCW12, PCW13, and PCW16)^21^ and mouse ovaries (E11.5, E12.5, E14.5, E16.5, and E18.5)^22^, selected to represent developmentally comparable time windows matched to the bovine samples **(Figure 1A).**

**Figure 1.**
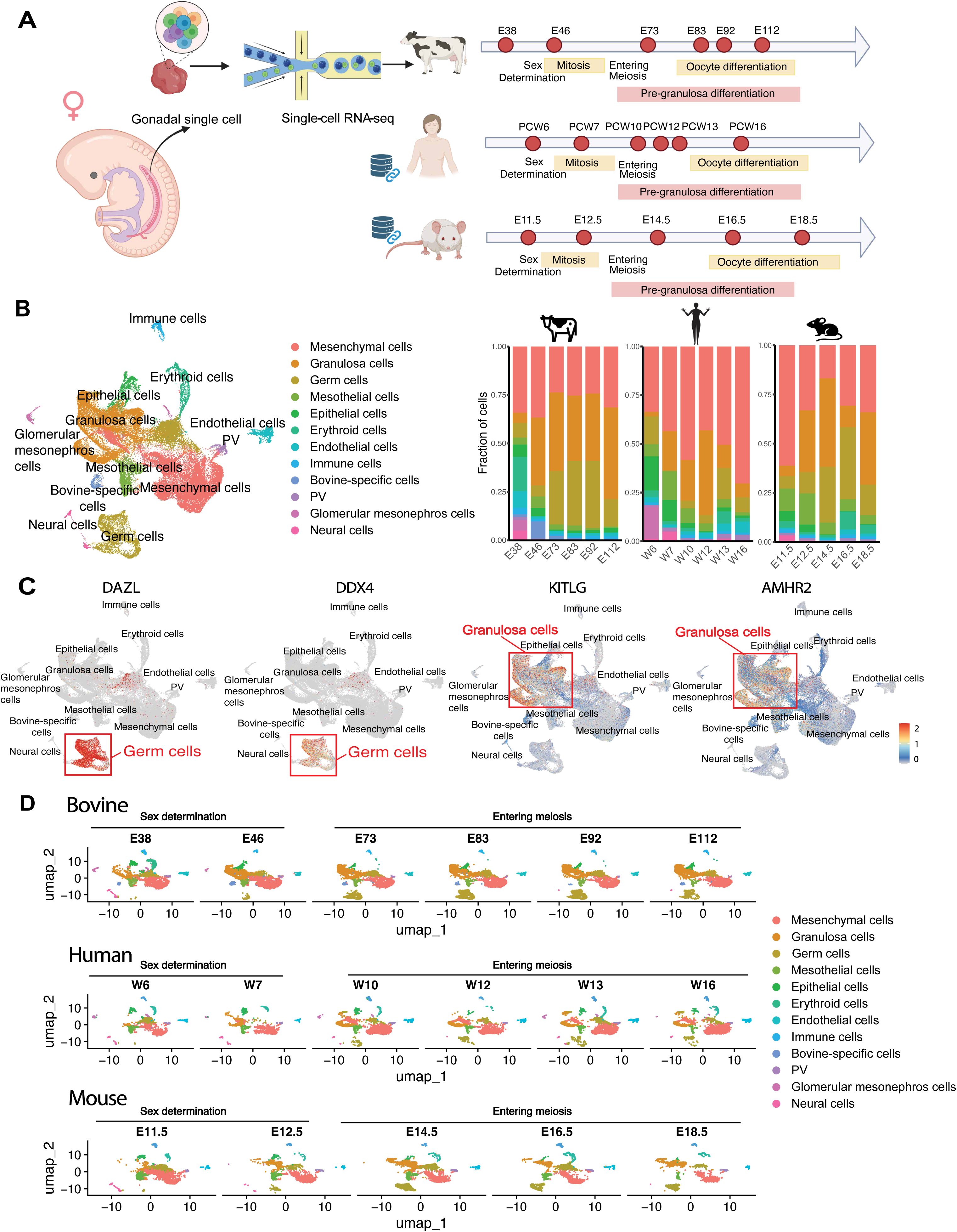
Cross-species single-cell transcriptomic atlas of female fetal gonadal development. **(A)** Schematic overview of the study design and developmental stages analyzed. Single-cell RNA sequencing was performed on bovine fetal gonadal tissues. Human and mouse fetal gonadal single-cell RNA-seq datasets were obtained from publicly available databases. Samples span key developmental stages including sex determination, mitotic expansion, entry into meiosis, and early oocyte differentiation. Corresponding embryonic or post-conception stages are indicated for each species. **(B)** UMAP showing clustering of major cell types in the integrated dataset. Twelve cell types were identified in total, 11 shared across species and one specific to cattle. PV indicates perivascular cells. Bar plots show the relative proportion of major cell types across developmental time points for each species, illustrating dynamic changes in cellular composition during ovarian development. **(C)** Feature plots showing the expression of canonical germ cell and granulosa cell markers projected onto the integrated UMAP. *DAZL* and *DDX4* mark germ cells, while *KITLG* and *AMHR2* mark granulosa cells. **(D)** UMAPs showing cell population distributions across three species and developmental time points.

Following integration based on 16,073 one-to-one orthologous genes, the combined dataset comprised 140,441 single cells across all three species **(Supplementary Table 1),** including 52,928 bovine, 48,874 human, and 38,639 mouse cells. After quality control, 107,930 high-quality cells were retained for downstream analysis **(Supplementary Table 1)**. Unsupervised clustering followed by manual annotation guided by marker genes established in human and mouse gonadal development^21^ identified 11 major cell types **(Figure 1B, Supplementary Table 2)**, including germ cells, granulosa cells, mesenchymal cells, mesothelial cells, epithelial cells, erythroid cells, endothelial cells, immune cells, perivascular (PV) cells, glomerular mesonephros cells (transient embryonic kidney), neural cells, as well as a population of unclassified bovine-specific cells. For example, germ cells were identified by high expression of *DAZL* and *DDX4* **(Figure 1C)**, granulosa cells by *KITLG* and *AMHR2* **(Figure 1C)**, epithelial cells by *PAX8*, and mesenchymal cells by *DCN* **(Supplementary Figure 1)**.

Across all species and developmental stages, mesenchymal cells, and granulosa cells together constituted the majority of the cellular population in the early development, highlighting their central roles during early ovarian development **(Figure 1B and 1D).** Interestingly, female germ cells became more predominant after the initiation of the meiosis stage, especially in bovine (E73) and mouse (E14.5) datasets. Moreover, the unclassified bovine-specific cell population was detected at all sampled stages, with a higher relative abundance at early gestational time points (E38, E46, and E73), suggesting the presence of species-specific cellular states during early bovine gonadal development **(Figure 1B and 1D)**.

### Dynamics of epigenetic regulators during early ovarian development

To investigate epigenetic reprogramming during early development of female germ cells and granulosa cells, we analyzed the gene expression dynamics of key regulators associated with DNA methylation (DNAme), histone modifications, and chromatin remodeling across the 3 species **(Figure 2; Supplementary Figure 2)**.

**Figure 2.**
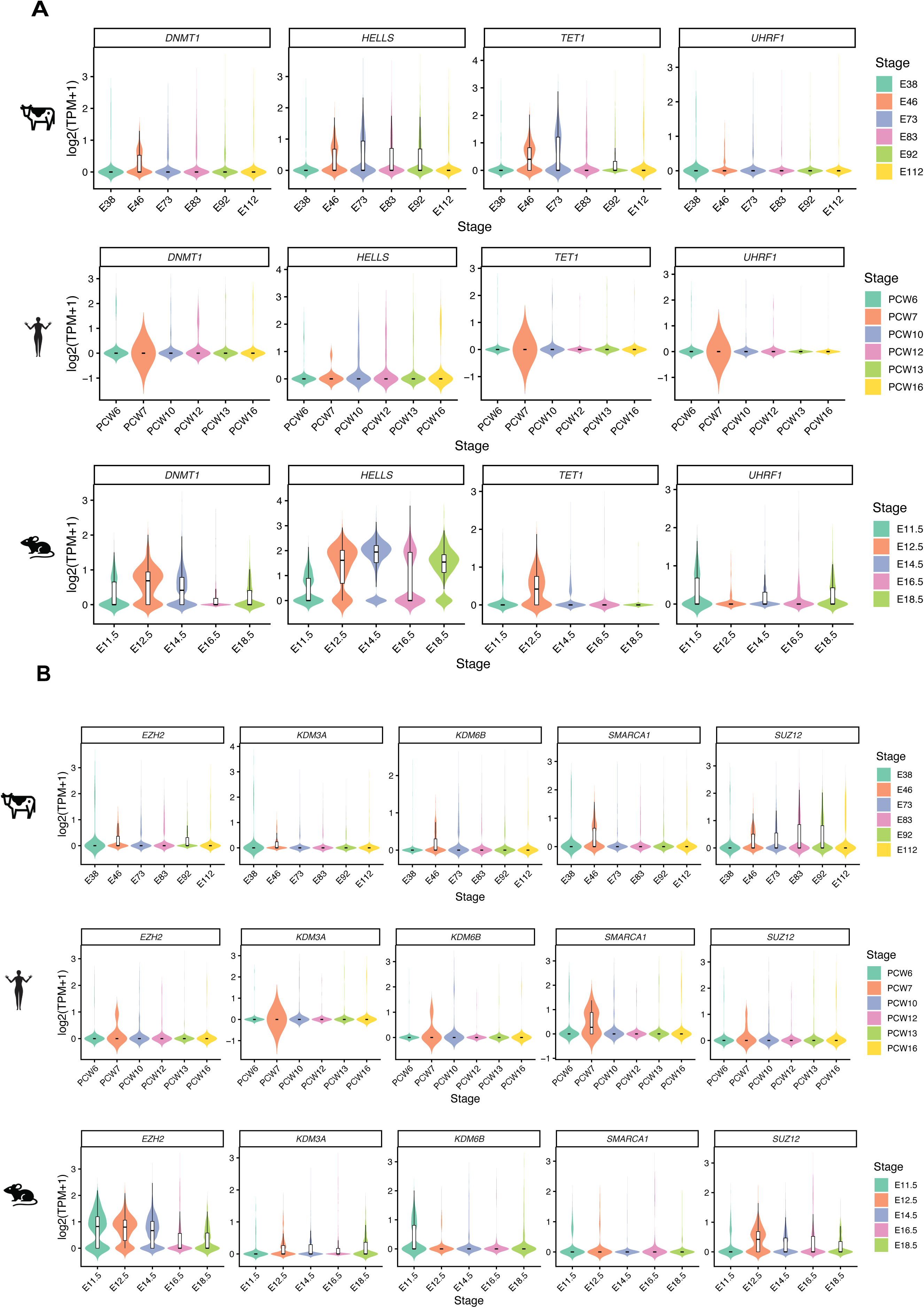
Cross-species dynamics of epigenetic regulators during early germ cell development. **(A)** Expression dynamics of DNA methylation–related regulators (*DNMT1*, *HELLS*, *TET1*, *UHRF1*) across developmental stages in bovine (E38-E112), human (PCW6–PCW16) and mouse (E11.5–E18.5) germ cells. **(B)** Expression dynamics of chromatin remodeling and histone modification–related regulators (*EZH2*, *KDM3A*, *KDM6B*, *SMARCA1*, *SUZ12*) across developmental stages in bovine (E38–E112), human (PCW6–PCW16), and mouse (E11.5–E18.5) germ cells. Violin plots show the distribution of log2(TPM + 1) expression levels for each gene at each developmental stage, with embedded boxplots indicating median and interquartile range. Colors represent developmental stages as indicated. Species are denoted by icons on the left (bovine, human, and mouse).

In female germ cells **(Figure 2)**, for DNAme regulators **(Figure 2A)**, we observed high expression of the maintenance DNA methyltransferase *DNMT1*^37^ and upregulation of the DNA demethylation factor *TET1*^38^ at early developmental stages (E46 in bovine; PCW7 in human and E12.5 in mouse) followed by decreased expression of *DNMT1* and *TET1* at later stages **(Figure 2A)**. This pattern is consistent with the well-characterized wave of global DNA demethylation followed by remethylation during germ cell development^39^.

Histone modification-related regulators also exhibited dynamic changes during early germ cell development **(Figure 2B)**. *EZH2*, a core component of the Polycomb repressive complex 2 (PRC2) responsible for H3K27 trimethylation^40^, showed sustained but gradually decreasing expression in mouse female germ cells (E11.5-E18.5). Histone demethylases *KDM3A*^41^ exhibited continuous activation in mouse germ cells after sex determination (E12.5-E18.5). Another histone demethylases *KDM3B*^41^ showed transient activation in bovine (E46) and mouse (E11.5). *SMARCA1*, a chromatin remodeler^42^, exhibited transient upregulation at early stages in bovine (E46) and human (PCW7), but not in mouse. In addition, *SUZ12*, another core component of the PRC2 complex required for its stability and catalytic activity^43^, was upregulated in bovine germ cells (E46-E83) but showed decreasing expression in mouse female germ cells (E12.5-E18.5).

In granulosa cells, for DNAme regulators **(Supplementary Figure 2A)**, expression of *DNMT1*^37^ showed a progressive decrease following sex determination in bovine (E46 onward) and human (PCW7 onward). In contrast, mouse *DNMT1* showed transient activation at E12.5 and E16.5. *HELLS*, a chromatin remodeler involved in DNA methylation maintenance and heterochromatin organization^44^, showed transient activation at E46 in bovine and at E11.5, E12.5 and E16.5 in mouse. In contrast, *TET1* displayed a gradual increase in expression after E73 in bovine, whereas in human and mouse it showed only transient upregulation at PCW13 and E16.5, respectively. *UHRF1*, another DNA methylation maintenance factor^45^, showed transient activation at E11.5 in mouse. Together, these patterns suggest a shift from DNA methylation maintenance toward increased demethylation activity during granulosa cell development, although this shift varies across species in both timing and extent.

Histone modification-related regulators such as *EZH2* showed dynamics similar to *DNMT1* in each species, indicating the consistency in methylome erasure and reestablishment during development **(Supplementary Figure 2B)**. In contrast, *KDM3A* and *KDM6B* showed distinct species-specific dynamics. In bovine, both genes were highly expressed at early stages (E38 and E46) and subsequently declined, whereas in human their expression was high at early development, showed a transient decrease after PCW12 and increased afterwards. In mouse, both *KDM3A* and *KDM6B* showed a sustained increase from E12.5 to E18.5. A similar divergence was observed for *SMARCA1*, which was highly expressed at early stages and gradually decreased in bovine, remained consistently high in human, and showed an increased expression pattern in mouse. Moreover, *SUZ12* showed broader activation windows across species, which sustained high expression in bovine, a decrease then increase dynamic pattern in human, and a pronounced peaked expression at E16.5 in mouse. Together, these results suggest that epigenetic changes may also occur in granulosa cells during early development, although it has been more extensively characterized in germ cells.

### Conserved and species-specific transcriptional dynamics during early germ cell development

To reconstruct the developmental progression of early germ cells, we performed pseudotime trajectory analysis of germ cells for each species **(Figure 3A; Supplementary Figure 3)**. Across all three species, germ cell development followed a consistent transition from pluripotency-associated states toward meiotic initiation and differentiation. A set of genes has been identified that exhibited conserved dynamic expression patterns during this process (**Figure 3A-B**). For example, early development was marked by high expression of pluripotency and germline specification factors, including *POU5F1*^46^, *NANOG*^47^, *TFAP2C*^48^, and *SOX17*^49^, consistent with an undifferentiated germ cell identity. As the development progressed, these markers were gradually downregulated, coinciding with the activation of germ cell maturation regulators such as *KIT*^50^, and RNA-binding factors *ELAVL2*^51^. At later stages, meiotic entry and germ cell differentiation were characterized by the upregulation of meiotic genes, including *SYCP1*^52^, *SYCP2*^53^, *SYCP3*^54^, *TEX30*^55^, *ZCWPW1*^56^, and *SMC1B*^57^ **(Figure 3A; supplementary Figure 3B)**. We then overlapped trajectory-associated genes across the three species and identified a total of 34 conserved, highly dynamic genes, including *NANOG, POU5F1*, *SYCP1*, and *JUN* **(Figure 3B)**. Beyond these conserved regulators, genes involved in cell-cycle control such as *UBE2C*^58^ and *TOP2A*^59^, translational regulation such as *CPEB1*^60^, and broader differentiation and signaling processes represented by *JUN*^61^ and *NR6A1*^62^, also displayed dynamic expression changes along pseudotime across all three species. In addition, a substantial number of genes displayed species-specific expression dynamics, including 162 bovine-specific, 36 human-specific, and 566 mouse-specific genes **(Figure 3B).** These differences indicate species-specific divergence in gene regulatory features despite a conserved core trajectory of germ cell development. For example, GO analysis of trajectory-associated genes in each species revealed species-specific pathway enrichment, with bovine female germ cells enriched for pathways related to cell morphogenesis, reproductive process, and neurodevelopment, human female germ cells enriched for cell division, mitotic cell cycle, and DNA replication processes, and mouse female germ cells enriched for meiosis-related pathways, including homologous chromosome pairing and segregation and meiotic cell cycle **(Figure 3C)**. These differences likely reflect variation in developmental timing and sampling across species.

**Figure 3.**
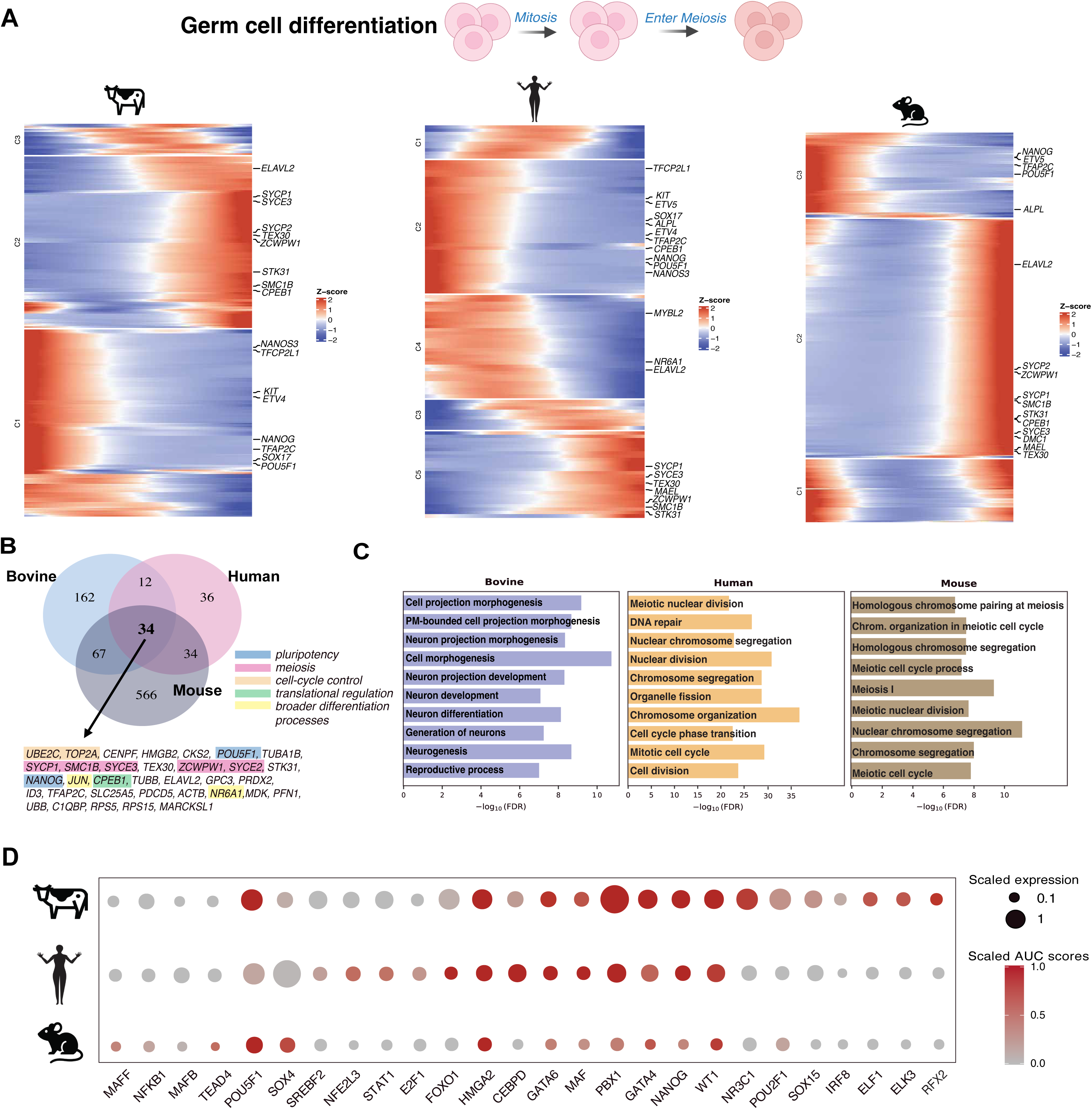
Conserved and species-specific transcriptional dynamics during early germ cell development across species. **(A)** Heatmaps showing gene expression dynamics along pseudotime in germ cells from bovine, human, and mouse. Cells are ordered from early to late pseudotime, and gene expression is scaled by gene (z-score). Pluripotency-associated genes and meiosis-related genes are highlighted, illustrating a progressive downregulation of pluripotency programs and activation of meiosis-associated transcriptional programs during germ cell development across species. **(B)** Venn diagram summarizing the overlap of dynamically expressed genes identified along pseudotime in bovine, human, and mouse germ cells**. (C)** Gene Ontology (GO) enrichment analysis of species-specific dynamic genes identified along pseudotime. **(D)** Transcription factor activity analysis inferred by pySCENIC. Dot size represents scaled expression levels, and color intensity indicates scaled regulon activity (AUC scores).

To further characterize regulatory programs during germ cell differentiation, we defined transcription factor regulon activity in each species **(Figure 3D)**. This analysis identified a set of conserved active regulons across species, including *NANOG*, *WT1*, *GATA4*, *GATA6*, *PBX1*, *MAF*, and *HMGA2*. Several of these regulators have been previously implicated in male gonadal or germline-related contexts. For example, *PBX1*, which shows the strongest signal in cattle, has been reported as a master regulator of Leydig cell differentiation and steroidogenesis-related function in mice^63^. Other regulators include *WT1*, which is required for testis development^64^, *MAF*, which contributes to germline stem cell maintenance^65^, and *HMGA2*, which has been associated with postpubertal testicular germ cell tumour^66^. How these regulators contribute to early female gonadal development remains to be determined.

In addition, several regulons exhibited species-specific activity patterns. For example, we found the enriched *ELF1*, *ELK3*, and *RFX2* activity in bovine germ cells, elevated *MAFF* and *TEAD4* activity in mouse germ cells, and increased *NFE2L3*, *STAT1*, and *E2F1* activity in human germ cells. *RFX2* is also a key transcriptional regulator of mouse spermiogenesis^67^, although its potential role in early female germ cell development in bovine remains unclear. *STAT1* is likely to be responsible for initiating transcription network in human fetal germ cells^68^. *E2F1* has been shown to regulate germ cell apoptosis during the first wave of spermatogenesis in mouse^69^. Together, these results demonstrate that while mammalian germ cell development follows a broadly conserved trajectory across species, it is accompanied by both shared and species-specific gene expression programs and transcription factor regulons that reflect divergence in gene regulatory control.

### Conserved and species-specific transcriptional dynamics during early granulosa cell development

Similarly, we performed pseudotime trajectory analysis for granulosa cells to study their development separately for each species **(Figure 4A; Supplementary Figure 4)**. In bovine granulosa cells, early development was marked by WNT signaling pathway modulator *SFRP*^70^, cell-cell adhesion regulator *CDH4*^71^, and *DCN*, which has a structural role in ovarian extracellular matrix^72^. At later stages, granulosa cells showed activation of genes associated with endocrine maturation and follicular support, including *INHA*^73^, *CYP11A1*^74^, and *PAPPA*^75^. In human granulosa cells, early to intermediate stages showed expression of *GPC3*, which encodes a glypican found in the extracellular matrix^22^, and *MDK*, a growth factor that is thought to support postnatal follicle growth in human ovaries^76^. Differentiated granulosa cells were marked by activation of key granulosa cell markers, including *FOXL2*^77^ and *GJA1*^78^, alongside IGF pathway modulation through *IGFBP2*^79^ and retinoic acid metabolism via *RDH10*^80^. In mouse granulosa cells, early developmental stage was also characterized by expression of developmental and growth factor associated regulators such as *GPC3*, *MDK*, and *LHX9,* a LIM homeobox transcription factor implicated in gonadal development and ovarian function in mouse models^81^. Late developmental stage in mice showed expression of genes involved in key signaling pathways, such as *DHH*, a Hedgehog ligand expressed in granulosa cells with roles in follicle signaling and theca differentiation^82^, and granulosa cell maturation was marked by upregulation of *FOXL2*, similar to humans **(Figure 4A)**.

**Figure 4.**
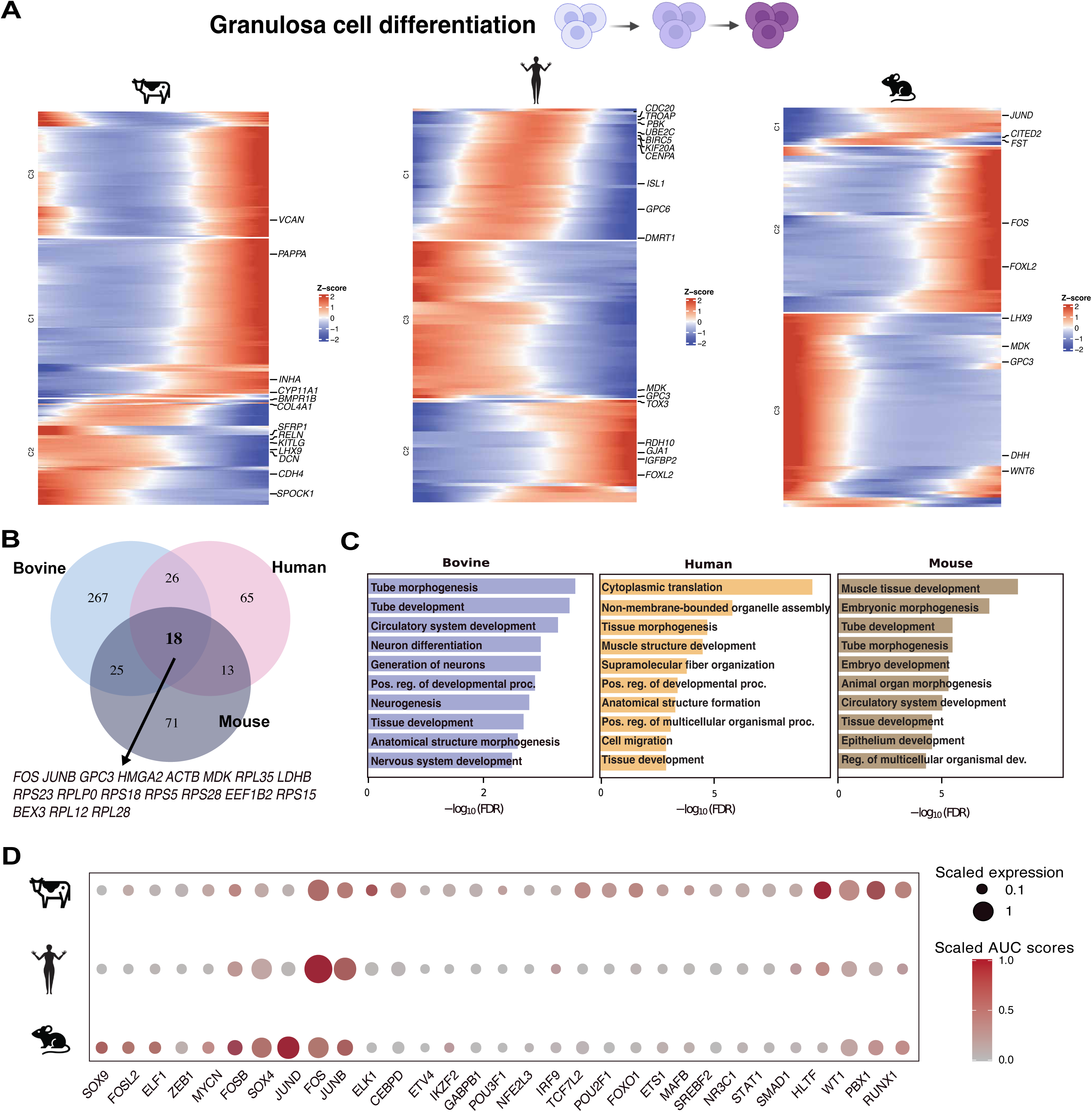
Conserved and species-specific transcriptional dynamics during early granulosa cell development across species. **(A)** Heatmaps showing gene expression dynamics along pseudotime in granulosa cells from bovine, human, and mouse. Cells are ordered from early to late pseudotime, and gene expression is scaled by gene (z-score). Representative genes associated with early supporting states, signaling pathways, and granulosa cell differentiation are highlighted, revealing progressive transcriptional changes during granulosa cell development across species. **(B)** Venn diagram summarizing the overlap of dynamically expressed genes identified along granulosa cell pseudotime in bovine, human, and mouse. **(C)** Gene Ontology (GO) enrichment analysis of species-specific dynamic genes identified along granulosa cell pseudotime. **(D)** Transcription factor activity analysis inferred by pySCENIC. Dot size represents scaled expression levels, and color intensity indicates scaled regulon activity (AUC scores).

Some of these genes were conserved across developmental stages in all three species, including *FOS* and *JUNB*, both members of the AP-1 transcription factor family^83^. These factors are induced in granulosa cells in response to gonadotropins and contribute to follicle maturation and ovulatory gene regulation in mice^84^ **(Figure 4B)**. Moreover, species-specific trajectory-associated genes were also observed with specific GO term enrichment, further indicating their regulatory divergence. Specifically, we observed that bovine granulosa cells were enriched for tube morphogenesis and neurogenesis, while human cells prioritized cytoplasmic translation, and mouse cells emphasized muscle tissue development **(Figure 4C)**.

Transcription factor regulon analysis also identified *FOS* and *JUNB* as the most enriched common active regulons across all three species **(Figure 4D)**, especially in humans. In contrast, several regulons exhibited species-specific activity patterns among granulosa cells. In bovine, *HLTF*, a DNA repair-associated factor^85^, and *PBX1*, a developmental transcriptional co-factor^86^, showed elevated activity. In mice, *JUND*^83^, a member of the AP-1 family, together with *FOS* and *JUNB*, exhibits increased regulon activity.

### Cell-cell interaction features between germ cells and gonadal somatic cells

To characterize how cell-cell communication among female gonadal somatic cell populations and early germ cells contributes to the gonadal differentiation and gametogenesis, we defined ligand-receptor interactions among cell types within each species **(Figure 5)**. Overall, interactions involving germ cells were relatively weak compared with those among gonadal somatic cell populations, which exhibited more frequent intercellular communications, a pattern consistently observed across all three species **(Figure 5A)**.

**Figure 5.**
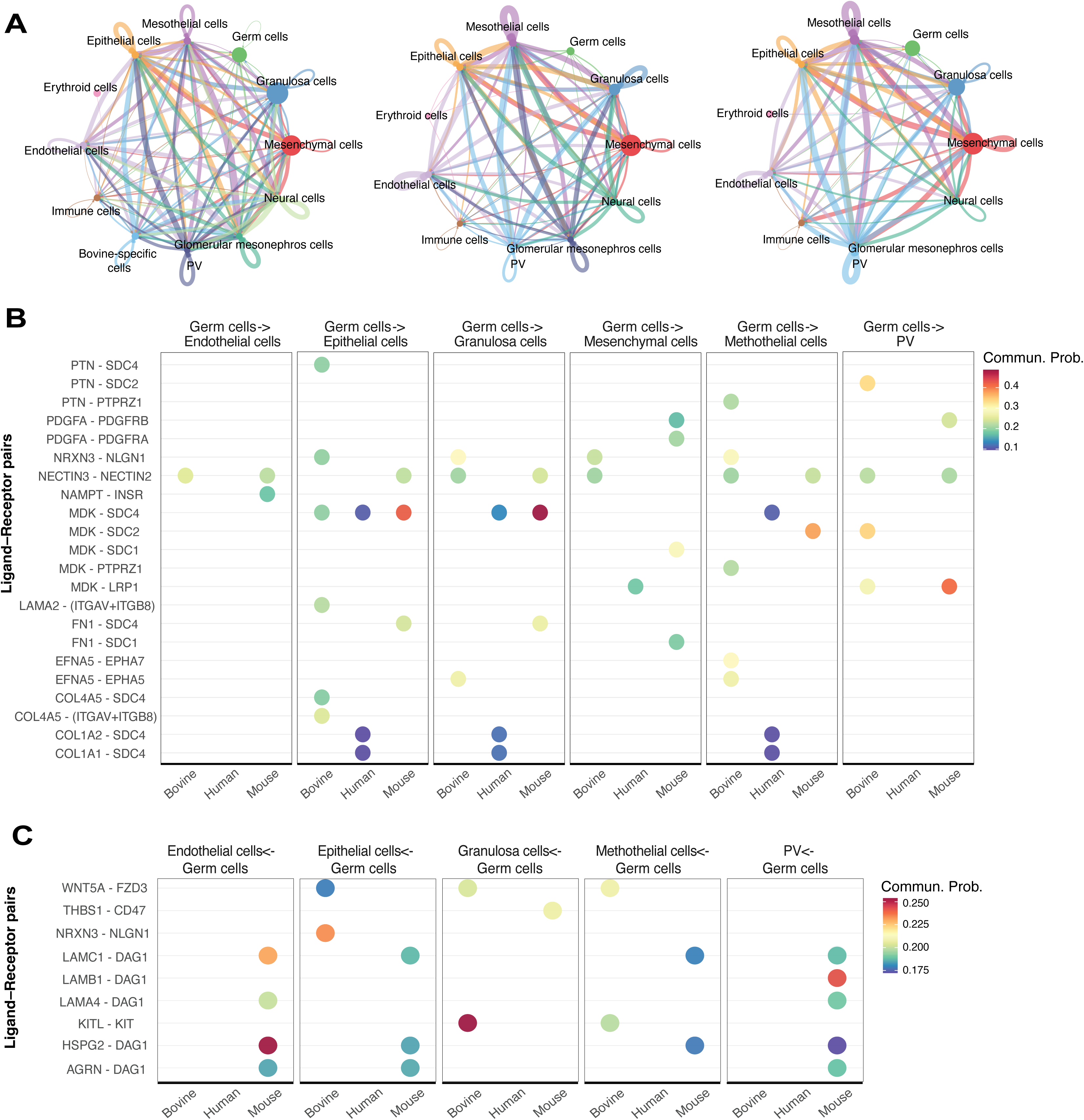
Cross-species cell–cell communication networks involving germ cells. **(A)** Global cell-cell communication networks inferred from single-cell transcriptomes of female fetal gonads in bovine (left), human (middle), and mouse (right). Nodes represent annotated cell types, and edges represent inferred ligand–receptor–mediated interactions. Edge thickness reflects the overall strength of communication between cell types. **(B)** Dot plots showing inferred ligand–receptor interactions with germ cells specified as the source cell population. Each panel summarizes outgoing signaling from germ cells to major somatic cell types across species. Dot size and color indicate the communication probability for each ligand–receptor pair. **(C)** Dot plots showing inferred ligand-receptor interactions with germ cells specified as the target cell population. Incoming signaling from surrounding somatic cell types, including endothelial, epithelial, granulosa, mesothelial, and PV cells, is shown across species. Dot size and color represent communication probability.

To further dissect germ cell-associated signaling, we first designated germ cells as the signaling source and examined ligand-receptor interactions between them and other gonadal cell types **(Figure 5B)**. Several ligand-receptor pairs were conserved across species. For instance, MDK-SDC4 interactions, involving the growth factor Midkine (MDK)^87^ and its co-receptor Syndecan-4 (SDC4)^88^, were detected between germ cells and epithelial cells in all three species and were additionally shared between germ cells and granulosa cells in both human and mouse. Moreover, NECTIN3-NECTIN2 interactions, which regulate cell-cell adhesion and junction formation^89^, were conserved between cattle and mouse and occurred between germ cells and multiple somatic cell types, including endothelial cells, granulosa cells, mesothelial cells, and perivascular cells (PV).

Species-specific ligand-receptor interaction pairs were also identified, for example, COL1A2-SDC4 and COLA1- SDC4 were only observed to signal out from human germ cells to either epithelial cells, granulosa cells, or mesothelial cells. EFNA5-EPHA5 were only sent from bovine germ cells into granulosa cells or mesothelial cells. FN1-SDC4 were only observed between mouse germ cells and epithelial cells or granulosa cells.

We next reversed the signaling direction by designating germ cells as the signal receptor to examine incoming signals from surrounding somatic cell types **(Figure 5C)**. In contrast to the analysis in which germ cells were treated as signaling sources, overall interaction strength was reduced, and only species-specific signaling pathways were identified. For example, WNT5A-FZD3 and KITL-KIT interactions, which are known to trigger PI3K/Akt signaling and promote fibroblast adhesion^90^, and to play crucial roles in primordial follicle formation, activation, and follicular development^91^, respectively, were observed between bovine granulosa cells and germ cells. THBS1-CD47 signaling was detected specifically between mouse granulosa cells and germ cells. Interestingly, we did not find significant enrichment of ligand-receptor interaction pairs between human somatic cells and germ cells.

### Cross-species cell type classification using a support vector machine (SVM) model

To identify conserved cell types across three species, we trained a support vector machine (SVM) classifier on the human dataset to distinguish major gonadal cell types and applied the trained model to predict mouse and bovine cell types (**Figure 6**). In mice, erythroid cells, immune cells, endothelial cells, neural cells, PV, and germ cells exhibited high prediction probabilities, close to 1, indicating strong cross-species conservation (**Figure 6A)**. Mesenchymal, mesothelial, and epithelial cells showed moderately high probabilities (∼0.6-0.8), suggesting partial conservation. In contrast, granulosa cells showed low prediction probabilities (<0.2), suggesting pronounced cross-species divergence on these cell types **(Figure 6A)**. In bovine, prediction patterns differed from those of the mouse (**Figure 6B)**. For example, glomerular mesonephros cells, endothelial cells, immune cells, neural cells, PV, erythroid cells, and germ cells showed high probabilities (>0.8). Mesothelial, mesenchymal, epithelial, and granulosa cells showed low prediction probabilities (<0.4), with granulosa cells being the lowest (**Figure 6B)**. These results indicate that some gonadal somatic cells such as immune cells exhibit more conserved transcriptomic profiles across species, as reflected by their higher cross-species prediction probabilities and greater overlap in UMAP space. In contrast, granulosa cells display clearer species-specific divergence, with distinct clustering observed across the three species **(Figure 6C).**

**Figure 6.**
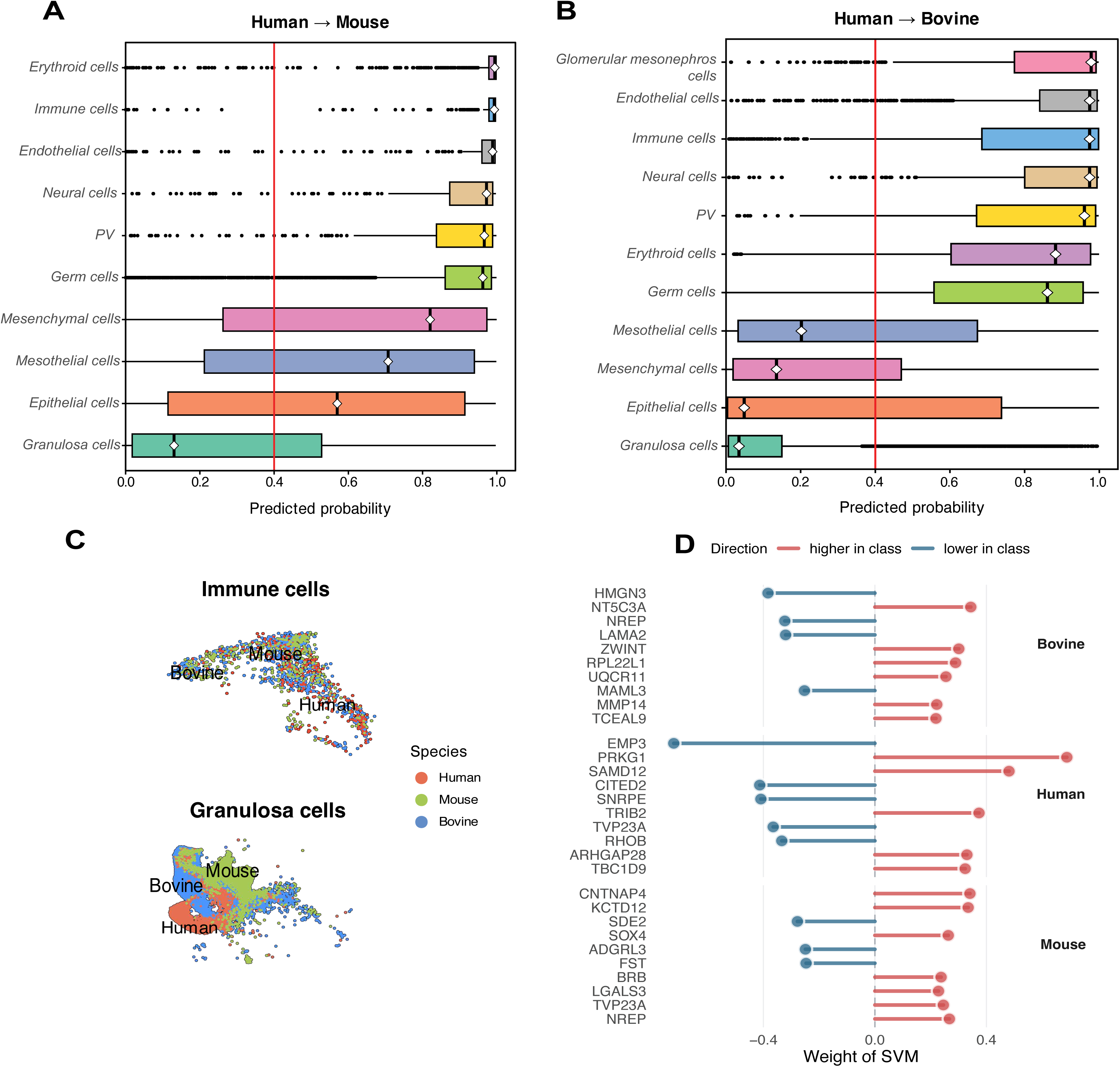
Cross-species comparison of cell types using a support vector machine (SVM)–based classifier. **(A)** Distribution of predicted classification probabilities for each cell type when a human-trained SVM classifier was applied to the mouse dataset. Each point represents an individual cell, and boxplots summarize the distribution of prediction probabilities for each annotated cell type. Higher prediction probabilities indicate greater transcriptional similarity to the corresponding human cell type. The vertical dashed red line indicates a reference probability threshold. **(B)** Distribution of predicted classification probabilities for each cell type when the human-trained SVM classifier was applied to the bovine dataset. Visualization and interpretation are as in panel **(A)**. **(C)** UMAP visualization of representative cell types illustrating cross-species similarity and divergence. Mesenchymal cells show substantial overlap across species, whereas granulosa cells display increased separation, consistent with reduced cross-species classification confidence. **(D)** Species-specific transcriptional features of granulosa cells identified by SVM. An additional SVM classifier was trained to distinguish granulosa cells from bovine, human, and mouse. The top weighted genes contributing to species-specific classification are shown, with positive and negative weights indicating relative enrichment in each species.

To further characterize which genes contribute to granulosa cell divergence across species, we trained a second SVM using granulosa cells from all three species and extracted genes with the highest model weights **(Figure 6D)**. In bovine granulosa cells, top-weighted genes, such as *LAMA2*^92^ and *MMP14*^93^, were associated with extracellular matrix organization and remodeling of the cellular microenvironment. In human granulosa cells, genes such as *RHOB*^94^ and *ARHGAP28*^95^ were enriched, implicating Rho GTPase-mediated cytoskeletal signaling. In mouse granulosa cells, genes including *SOX4*^96^ and *FST*^97^ were prominent, which were related to TGF-β/Activin-related signaling. Together, these features highlight species-specific regulatory features in granulosa cells.

Finally, to investigate the identity of the unclassified bovine-specific cell population **(Figure 1B)**, we trained an SVM on bovine data excluding this population and used the model to predict its closest cell type identity **(Supplementary Figure 5A).** The majority of cells were classified as mesenchymal or granulosa cells, together accounting for >90% of predictions, with smaller contributions from germ cells and epithelial cells. Compared with mesenchymal cells, the unclassified bovine-specific population showed a distinct pattern, such as increased expression of *FXYD3*^98^, involved in Na⁺/K⁺-ATPase–mediated ion regulation, and *HS3ST5*^99^, which functions in heparan sulfate sulfotransferase **(Supplementary Figure 5B)**. Indeed, this population did not express canonical granulosa cell markers but instead partially expressed the mesenchymal lineage-associated marker *PDGFRA*^100^ **(Supplementary Figure 5C)**. As early ovarian somatic cells include precursors of multiple lineages, including steroidogenic cells, we next assessed the expression of steroidogenesis-related genes. These cells showed clear expression of *CYP17A1*^101^, and partial expression of genes involved in steroid production and transport, including *INSL3*^102^*, HSB3D1*^103^, and STAR^104^ **(Supplementary Figure 5C)**. Together, these results suggest that the unclassified bovine-specific somatic population may exhibit molecular features consistent with a steroidogenic lineage.

## Discussion

We constructed an integrated single-cell transcriptomics atlas of early female gonadal development across bovine, human, and mouse to systematically identify conserved and divergent features of gonadal differentiation and define underlying molecular regulatory mechanisms. Overall, we identified 11 shared gonadal cell types, as well as a bovine-specific cell population that may exhibit steroidogenic features. Epigenetic regulators exhibited dynamic changes in both germ cells and granulosa cells. Developmental trajectory analysis revealed common transcriptional dynamic pattern regulating germ cell meiotic entry and granulosa cell differentiation, alongside divergence in the top functional gene pathways and regulons in each species. Cell-cell communication analysis further demonstrated that early gonad development is supported by extensive and dynamic signaling interactions between germ cells and surrounding somatic cells. Finally, implementation of a support vector machine (SVM)-based framework enabled quantitative assessment of transcriptional conservation and provided a predictive model for cross-species cell lineage classification.

### Classification of a bovine-specific cell type

In our analysis, we used an SVM-based classifier to identify a unique bovine-specific cell type, which characterized by steroidogenic features, including expression of *CYP17A1* and *HSD3B1*. In humans, *CYP17A1* expression was not detected in early female gonadal cell types between PCW5-12, however, it was observed in male Leydig and interstitial cells in the same time window^105^. *CYP17A1* is also a known marker of theca cells^106^, which typically differentiate in the ovarian follicle around puberty in females. In mice, theca cells progenitors arise from two populations, including an ovary-intrinsic precursor population detectable as early as E10.5 in the genital ridge, and a mesenchymal population originating from the mesonephros that migrates into the developing ovary around E12.5^107^. However, neither of these theca precursor populations exhibits strong steroidogenic gene expression (e.g., *HSD3B1*) during fetal development^107^. In contrast, we identified a bovine-specific cell population expressing steroidogenic genes (*CYP17A1* and *HSD3B1*) that presents from E38-E112, and does not align with the developmental patterns described in humans and mice. This divergence in both gene expression and developmental timing suggests that this population may represent either a bovine-specific early steroidogenic lineage in the female gonad or a theca precursor population with species-specific temporal state.

### Conserved and divergent regulatory programs in germ cell and granulosa cell development

Our developmental trajectory analysis shows a conserved gene expression pattern during the transition from PGCs to meiotic entry across cattle, humans, and mice. This process begins with the downregulation of pluripotency factors *POU5F1*^46^, *NANOG*^47^, and *SOX17*^49^, followed by the activation of maturation markers like *KIT*^50^ and *ELAVL2*^51^. While *SOX17* is known as a key specifier in humans^49^, its consistent dynamic in our bovine data confirms its potential role for ungulate germline development as well. All three species eventually activate the same meiotic program, including structural genes *SYCP1*^52^, *SYCP2*^53^, *SYCP3*^54^, and the epigenetic regulator *ZCWPW1*^56^. The shared timing of these genes across species suggests that the core mechanism for entering meiosis is highly stable in mammals.

Similarly, granulosa cell trajectories highlight a conserved role for the AP-1 family, specifically *FOS* and *JUNB*^83^. These factors act as common regulators for follicle maturation across all three species. While a previous study showed they coordinate metabolic shifts and cholesterol synthesis in adult periovulatory granulosa cells^108^, our results suggest they may also play a role in embryonic granulosa cells. Interestingly, *PBX1* showed the strongest regulon activity in bovine germ cells and was also elevated in bovine granulosa cells compared to humans and mice. Since *PBX1* is known to regulate genes involved in steroid production^63^, its high activity in both cell types may be linked to the bovine-specific cell population we identified that expresses steroidogenic markers. This suggests that in cattle, *PBX1* might help coordinate a specific endocrine environment during early gonadal development that is not as prominent in the other two species. *FOXL2* also showed species divergence. As a critical granulosa cell determinant in humans and mice^21^, it was detected at late pseudotime in these species but not in bovine, suggesting a delayed granulosa cell maturation program in cattle, in contrast to its strong expression in adult bovine follicular granulosa cells^24^.

### Cell-cell communication between germ cells and gonadal somatic cells

Cell-cell signaling interaction analysis further provides insight into how the germline niche may be established during early gonadal development. Compared with the extensive communication among somatic cell populations, germ cells-associated signaling was relatively limited, with germ cells primarily acting as signaling sources rather than receivers. This pattern raises the possibility that early germ cells may contribute to shaping their local microenvironment rather than functioning solely as passive recipients of niche signals. Conserved ligand-receptor pairs were observed, including MDK–SDC4 in all species and NECTIN3–NECTIN2 in cattle and mice, suggesting that growth factor signaling and adhesion-mediated interactions may represent conserved mechanisms contributing to germ cell niche establishment. *MDK* shows prominent expression in human granulosa cells, and MDK-syndecan interactions have been reported to regulate primordial follicle formation in humans and mice^109^. Similarly, nectin family adhesion molecules mediate cell–cell junctions in ovarian follicles^110^, and NECTIN2 has been observed to be expressed at cell-cell adhesion sites within the granulosa cell layer in the primary and preantral follicles in mice^111^, suggesting that NECTIN3-NECTIN2 interactions may play a conserved role in mediating cell-cell adhesion during early gonadal development in cattle and mice.

### Cross-species cell type classification and prediction

In addition, our cross-species cell type classification using an SVM revealed differential levels of transcriptional conservation among cell types. germ cells exhibit greater cross-species similarity, whereas granulosa cells display more pronounced cross-species divergence. This contrast likely reflects the fundamental and evolutionarily conserved roles of germ cells in early gonadal development^112^. In contrast, granulosa cells’ functions are more closely associated with species-specific follicular development and reproductive physiology^113^.

The top-weighted genes identified by the SVM classifier for species-specific granulosa cell discrimination converge on genes related to extracellular matrix (ECM) organization and microenvironmental regulation. In cattle, highly ranked genes such as *LAMA2*^92^ and *MMP14*^93^ are directly involved in ECM structure and remodeling. In humans, genes associated with Rho GTPase-mediated cytoskeletal signaling were enriched, a pathway closely linked to cell–matrix interactions and mechanical sensing^114^. In mice, the top genes are components of TGF-β/Activin signaling, which plays a key role in regulating ECM production and tissue remodeling^115^. Together, these results suggest that interspecies differences in granulosa cells are likely associated with divergence in ECM-related regulatory programs. Beyond revealing patterns of transcriptional conservation across species, SVM classifier also provides a predictive framework for cross-species cell type identification^116^. For example, by leveraging conserved transcriptional signatures, SVM classifier perform the best among 22 single cell classifiers in annotating cell populations in newly generated single-cell datasets, particularly in cases where canonical marker genes are incomplete, poorly characterized, or absent^116^. Such a strategy may be especially valuable for studies in non-model organisms, such as ruminant livestock, where cell type annotation often remains challenging^117^. Moreover, SVM classifier offers a scalable framework for integrating comparative single-cell datasets and for systematically identifying conserved and divergent cellular programs across species.

Despite these insights, several limitations of cross-species transcriptomic analysis should be considered. First, cross-species comparisons rely on approximate developmental stage alignment, and differences in sampling windows and dataset composition may influence the detection of conserved or species-specific cell populations. Second, our analysis is restricted to one-to-one orthologous genes, which limits the ability to capture species-specific genes or gene family expansions that may play important roles in lineage diversification. Future studies could address these limitations by expanding gene mapping strategies by incorporating all ortholog types and by integrating additional layers of information, such as regulatory or epigenetic features, to improve the robustness of cross-species comparisons.

## Conclusion

In summary, our integrative cross-species single-cell analysis provides a comprehensive framework for understanding early ovarian development across mammals. We identified both conserved gene regulatory programs and species-specific features that shape female germline and gonadal somatic cell differentiation. Our findings reveal a high degree of conservation in the transcriptional programs underlying germ cell development and meiotic entry, supporting the conservation of core developmental mechanisms. In contrast, granulosa cells exhibit species-specific divergent extracellular matrix-related gene regulatory networks, indicating how evolutionary differences may shape reproductive strategies and developmental timing. Beyond these biological insights, our study establishes a generalizable framework for cross-species single-cell transcriptomic integration, quantitative assessment of transcriptional conservation, and cell type annotation. This resource lays the groundwork for future comparative studies in reproductive biology and offers a foundation for exploring cell-type-specific regulatory programs across species.

## Declarations

### Ethics approval and consent to participate

All animal experimental procedures were performed in accordance with the National Research Council’s Guide for the Care and Use of Laboratory Animals and were approved by the Institutional Animal Care and Use Committee (IACUC) of Northwest A&F University (protocol code: IACUC2024-1204, date of approval: 10 December 2024).

### Data and Resource Availability

RNA-seq data were deposited in the China National Center for Bioinformation (CNCB) under accession code: CNCB: OMIX012725. Human and mouse single-cell RNA-seq datasets used in this study are publicly accessible from ArrayExpress (E-MTAB-10551) and GEO (GSE136441).

### Authors’ contributions

Y.F. performed data processing, integration, computational analyses, and figure preparation. S.H. collected the samples and conducted the experiments. J.Z., S.X., and Y.S. assisted in sample collection. J.E.D., T.Y., and Y.W. supervised the study and provided conceptual guidance. Y.F. drafted the manuscript, and all authors reviewed, revised, and approved the final version.

## Acknowledgements

The authors acknowledge all members from the Tang lab and Wei lab for assistance with sample collection, and all members from the Duan lab for support with study design and data analysis.

## References

1. Mamsen LS, Brøchner CB, Byskov AG, Møllgard K. The migration and loss of human primordial germ stem cells from the hind gut epithelium towards the gonadal ridge. The International Journal of Developmental Biology. 2013;56(10-11-12):771-778. doi:10.1387/ijdb.120202lm

2. Anderson R, Copeland TK, Schöler H, Heasman J, Wylie C. The onset of germ cell migration in the mouse embryo. Mechanisms of Development. 2000;91(1):61–68. doi:10.1016/S0925-4773(99)00271-3

3. Guiltinan C, Botigelli RC, Arcanjo RB, et al. Molecular profiling of bovine primordial germ cell specification and migration onset reveals a conserved program in bilaminar disc embryos. bioRxiv. Preprint posted online June 8, 2025:2025.06.04.657951. doi:10.1101/2025.06.04.657951

4. Tanaka SS, Nishinakamura R. Regulation of male sex determination: genital ridge formation and Sry activation in mice. Cell Mol Life Sci. 2014;71(24):4781–4802. doi:10.1007/s00018-014-1703-3

5. Xie Y, Wu C, Li Z, Wu Z, Hong L. Early Gonadal Development and Sex Determination in Mammal. Int J Mol Sci. 2022;23(14):7500. doi:10.3390/ijms23147500

6. Piprek RP. Genetic mechanisms underlying male sex determination in mammals. J Appl Genet. 2009;50(4):347–360. doi:10.1007/BF03195693

7. Chassot AA, Ranc F, Gregoire EP, et al. Activation of β-catenin signaling by Rspo1 controls differentiation of the mammalian ovary. Hum Mol Genet. 2008;17(9):1264–1277. doi:10.1093/hmg/ddn016

8. Borum K. Oogenesis in the mouse: A study of the meiotic prophase. Experimental Cell Research. 1961;24(3):495–507. doi:10.1016/0014-4827(61)90449-9

9. Lee SM, Surani MA. Epigenetic reprogramming in mouse and human primordial germ cells. Exp Mol Med. 2024;56(12):2578–2587. doi:10.1038/s12276-024-01359-z

10. Gill ME, Hu YC, Lin Y, Page DC. Licensing of gametogenesis, dependent on RNA binding protein DAZL, as a gateway to sexual differentiation of fetal germ cells. Proceedings of the National Academy of Sciences. 2011;108(18):7443–7448. doi:10.1073/pnas.1104501108

11. Yan A, Xiong J, Zhu J, et al. DAZL regulates proliferation of human primordial germ cells by direct binding to precursor miRNAs and enhances DICER processing activity. Nucleic Acids Res. 2022;50(19):11255–11272. doi:10.1093/nar/gkac856

12. Hickford DE, Frankenberg S, Pask AJ, Shaw G, Renfree MB. DDX4 (VASA) Is Conserved in Germ Cell Development in Marsupials and Monotremes1. Biol Reprod. 2011;85(4):733–743. doi:10.1095/biolreprod.111.091629

13. Xu C, Cao Y, Bao J. Building RNA-protein germ granules: insights from the multifaceted functions of DEAD-box helicase Vasa/Ddx4 in germline development. Cell Mol Life Sci. 2021;79(1):4. doi:10.1007/s00018-021-04069-1

14. Anderson EL, Baltus AE, Roepers-Gajadien HL, et al. Stra8 and its inducer, retinoic acid, regulate meiotic initiation in both spermatogenesis and oogenesis in mice. Proc Natl Acad Sci U S A. 2008;105(39):14976–14980. doi:10.1073/pnas.0807297105

15. Le Bouffant R, Guerquin MJ, Duquenne C, et al. Meiosis initiation in the human ovary requires intrinsic retinoic acid synthesis. Hum Reprod. 2010;25(10):2579–2590. doi:10.1093/humrep/deq195

16. Tachibana-Konwalski K, Godwin J, Weyden L van der, et al. Rec8-containing cohesin maintains bivalents without turnover during the growing phase of mouse oocytes. Genes Dev. 2010;24(22):2505–2516. doi:10.1101/gad.605910

17. Garcia-Cruz R, Brieño MA, Roig I, et al. Dynamics of cohesin proteins REC8, STAG3, SMC1β and SMC3 are consistent with a role in sister chromatid cohesion during meiosis in human oocytes. Hum Reprod. 2010;25(9):2316–2327. doi:10.1093/humrep/deq180

18. Global Disorders of Sex Development Update since 2006: Perceptions, Approach and Care | Hormone Research in Paediatrics | Karger Publishers. Accessed November 23, 2025. https://karger.com/hrp/article-abstract/85/3/158/166041/Global-Disorders-of-Sex-Development-Update-since?redirectedFrom=fulltext&casa_token=NS6Y0mhFVJsAAAAA:9_UBSb6PBzrS1F7gm5rvp6ME7KHUUQJpI56SCv3wHuMAwHf6tbqCEzEuEj0n-je8OfM-1x47

19. Hersmus R, de Leeuw BH, Stoop H, et al. A novel SRY missense mutation affecting nuclear import in a 46,XY female patient with bilateral gonadoblastoma. Eur J Hum Genet. 2009;17(12):1642–1649. doi:10.1038/ejhg.2009.96

20. Settergren I. Ovarian hypoplasia in heifers due to germ cell weakness. Theriogenology. 1997;47(2):531–539. doi:10.1016/S0093-691X(97)00011-3

21. Garcia-Alonso L, Lorenzi V, Mazzeo CI, et al. Single-cell roadmap of human gonadal development. Nature. 2022;607(7919):540–547. doi:10.1038/s41586-022-04918-4

22. Niu W, Spradling AC. Two distinct pathways of pregranulosa cell differentiation support follicle formation in the mouse ovary. Proceedings of the National Academy of Sciences. 2020;117(33):20015–20026. doi:10.1073/pnas.2005570117

23. Chen M, Long X, Chen M, et al. Integration of single-cell transcriptome and chromatin accessibility of early gonads development among goats, pigs, macaques, and humans. Cell Reports. 2022;41(5):111587. doi:10.1016/j.celrep.2022.111587

24. Han S, Zou Q, Fang Y, et al. Integration of Single-cell Transcriptome of Female Early Gonadal Development in Bovine, Goats, and Pigs. iScience, under publication process.

25. Zheng GXY, Terry JM, Belgrader P, et al. Massively parallel digital transcriptional profiling of single cells. Nat Commun. 2017;8(1):14049. doi:10.1038/ncomms14049

26. Hao Y, Stuart T, Kowalski MH, et al. Dictionary learning for integrative, multimodal and scalable single-cell analysis. Nat Biotechnol. 2024;42(2):293–304. doi:10.1038/s41587-023-01767-y

27. Germain PL, Lun A, Garcia Meixide C, Macnair W, Robinson MD. Doublet identification in single-cell sequencing data using scDblFinder. F1000Res. 2022;10:979. doi:10.12688/f1000research.73600.2

28. Zappia L, Oshlack A. Clustering trees: a visualization for evaluating clusterings at multiple resolutions. Gigascience. 2018;7(7):giy083. doi:10.1093/gigascience/giy083

29. Van den Berge K, Roux de Bézieux H, Street K, et al. Trajectory-based differential expression analysis for single-cell sequencing data. Nat Commun. 2020;11(1):1201. doi:10.1038/s41467-020-14766-3

30. Zhang J, Li H, Tao W, Zhou J. ClusterGVis: An Advanced Visualization and Clustering Tool for Gene Expression Analysis. genom proteom bioinform. Published online January 22, 2026:qzag005. doi:10.1093/gpbjnl/qzag005

31. Ge SX, Jung D, Yao R. ShinyGO: a graphical gene-set enrichment tool for animals and plants. Bioinformatics. 2020;36(8):2628–2629. doi:10.1093/bioinformatics/btz931

32. Kumar N, Mishra B, Athar M, Mukhtar S. Inference of Gene Regulatory Network from Single-Cell Transcriptomic Data Using pySCENIC. In: Mukhtar S, ed. Modeling Transcriptional Regulation: Methods and Protocols. Springer US; 2021:171–182. doi:10.1007/978-1-0716-1534-8_10

33. Jin S, Guerrero-Juarez CF, Zhang L, et al. Inference and analysis of cell-cell communication using CellChat. Nat Commun. 2021;12(1):1088. doi:10.1038/s41467-021-21246-9

34. Cortes C, Vapnik V. Support-vector networks. Mach Learn. 1995;20(3):273–297. doi:10.1007/BF00994018

35. Chen Y, Chen L, Lun ATL, Baldoni PL, Smyth GK. edgeR v4: powerful differential analysis of sequencing data with expanded functionality and improved support for small counts and larger datasets. Nucleic Acids Res. 2025;53(2):gkaf018. doi:10.1093/nar/gkaf018

36. Fan RE, Chang KW, Hsieh CJ, Wang XR, Lin CJ. LIBLINEAR: A Library for Large Linear Classification. Journal of Machine Learning Research. 2008;9(61):1871–1874.

37. Svedružić ŽM. Dnmt1 structure and function. Prog Mol Biol Transl Sci. 2011;101:221–254. doi:10.1016/B978-0-12-387685-0.00006-8

38. Jin C, Lu Y, Jelinek J, et al. TET1 is a maintenance DNA demethylase that prevents methylation spreading in differentiated cells. Nucleic Acids Res. 2014;42(11):6956–6971. doi:10.1093/nar/gku372

39. Guibert S, Forné T, Weber M. Global profiling of DNA methylation erasure in mouse primordial germ cells. Genome Res. 2012;22(4):633–641. doi:10.1101/gr.130997.111

40. Xie W, Chu Q, Brea L, et al. EZH2 PROTACs outperform catalytic inhibitors in prostate cancer by targeting a methylation-independent function of PRC2. Oncogene. 2026;45(5):636–649. doi:10.1038/s41388-025-03662-z

41. Macedo-Silva C, Miranda-Gonçalves V, Lameirinhas A, et al. JmjC-KDMs KDM3A and KDM6B modulate radioresistance under hypoxic conditions in esophageal squamous cell carcinoma. Cell Death Dis. 2020;11(12):1068. doi:10.1038/s41419-020-03279-y

42. Patil PA, Lombardo K, Sturtevant A, Mangray S, Yakirevich E. Loss of Expression of a Novel Chromatin Remodeler SMARCA1 in Soft Tissue Sarcoma. J Cytol Histol. 2018;9(6):524. doi:10.4172/2157-7099.1000524

43. Gong L, Liu X, Jiao L, Yang X, Lemoff A, Liu X. CK2-mediated phosphorylation of SUZ12 promotes PRC2 function by stabilizing enzyme active site. Nat Commun. 2022;13(1):6781. doi:10.1038/s41467-022-34431-1

44. Cousu C, Mulot E, De Smet A, et al. Germinal center output is sustained by HELLS-dependent DNA-methylation-maintenance in B cells. Nat Commun. 2023;14(1):5695. doi:10.1038/s41467-023-41317-3

45. Kong X, Chen J, Xie W, et al. Defining UHRF1 Domains That Support Maintenance of Human Colon Cancer DNA Methylation and Oncogenic Properties. Cancer Cell. 2019;35(4):633–648.e7. doi:10.1016/j.ccell.2019.03.003

46. Looijenga LHJ, Stoop H, de Leeuw HPJC, et al. POU5F1 (OCT3/4) identifies cells with pluripotent potential in human germ cell tumors. Cancer Res. 2003;63(9):2244–2250.

47. Chambers I, Silva J, Colby D, et al. Nanog safeguards pluripotency and mediates germline development. Nature. 2007;450(7173):1230–1234. doi:10.1038/nature06403

48. Chen D, Liu W, Zimmerman J, et al. The TFAP2C-Regulated *OCT4* Naive Enhancer Is Involved in Human Germline Formation. Cell Reports. 2018;25(13):3591–3602.e5. doi:10.1016/j.celrep.2018.12.011

49. Irie N, Weinberger L, Tang WWC, et al. SOX17 Is a Critical Specifier of Human Primordial Germ Cell Fate. Cell. 2015;160(1-2):253–268. doi:10.1016/j.cell.2014.12.013

50. Rossi P. Transcriptional control of KIT gene expression during germ cell development. Int J Dev Biol. 2013;57(2-4):179–184. doi:10.1387/ijdb.130014pr

51. Wiszniak SE, Dredge BK, Jensen KB. HuB (elavl2) mRNA is restricted to the germ cells by post-transcriptional mechanisms including stabilisation of the message by DAZL. PLoS One. 2011;6(6):e20773. doi:10.1371/journal.pone.0020773

52. Dunce JM, Dunne OM, Ratcliff M, et al. Structural basis of meiotic chromosome synapsis through SYCP1 self-assembly. Nat Struct Mol Biol. 2018;25(7):557–569. doi:10.1038/s41594-018-0078-9

53. Feng J, Fu S, Cao X, et al. Synaptonemal complex protein 2 (SYCP2) mediates the association of the centromere with the synaptonemal complex. Protein Cell. 2017;8(7):538–543. doi:10.1007/s13238-016-0354-6

54. Syrjänen JL, Pellegrini L, Davies OR. A molecular model for the role of SYCP3 in meiotic chromosome organisation. eLife. 2014;3:e02963. doi:10.7554/eLife.02963

55. Sun Q, Li Z, Wang B, et al. Loss of TEX30 leads to male subfertility in mice. Biol Reprod. 2025;113(3):672–685. doi:10.1093/biolre/ioaf163

56. Wells D, Bitoun E, Moralli D, et al. ZCWPW1 is recruited to recombination hotspots by PRDM9 and is essential for meiotic double strand break repair. de Massy B, Tyler JK, de Massy B, eds. eLife. 2020;9:e53392. doi:10.7554/eLife.53392

57. Islam KN, Modi MM, Siegfried KR. The Zebrafish Meiotic Cohesin Complex Protein Smc1b Is Required for Key Events in Meiotic Prophase I. Front Cell Dev Biol. 2021;9. doi:10.3389/fcell.2021.714245

58. Nicolau-Neto P, Palumbo A, De Martino M, et al. UBE2C Is a Transcriptional Target of the Cell Cycle Regulator FOXM1. Genes (Basel). 2018;9(4):188. doi:10.3390/genes9040188

59. Arroyo M, Fernández-Mimbrera MA, Gollini E, Esteve-Codina A, Sánchez A, Marchal JA. TOP2A inhibition and its cellular effects related to cell cycle checkpoint adaptation pathway. Sci Rep. 2025;15(1):3831. doi:10.1038/s41598-025-87895-8

60. Gebauer F, Hentze MW. Fertility Facts: Male and Female Germ Cell Development Requires Translational Control by CPEB. Molecular Cell. 2001;8(2):247–249. doi:10.1016/S1097-2765(01)00326-4

61. Raivich G, Behrens A. Role of the AP-1 transcription factor c-Jun in developing, adult and injured brain. Prog Neurobiol. 2006;78(6):347–363. doi:10.1016/j.pneurobio.2006.03.006

62. Lan ZJ, Xu X, Chung ACK, Cooney AJ. Extra-Germ Cell Expression of Mouse Nuclear Receptor Subfamily 6, Group A, Member 1 (NR6A1)1. Biol Reprod. 2009;80(5):905–912. doi:10.1095/biolreprod.107.067322

63. Wang FC, Zhang XN, Wu SX, He Z, Zhang LY, Yang QE. Loss of PBX1 function in Leydig cells causes testicular dysgenesis and male sterility. Cell Mol Life Sci. 2024;81(1):212. doi:10.1007/s00018-024-05249-5

64. Gao F, Maiti S, Alam N, et al. The Wilms tumor gene, Wt1, is required for Sox9 expression and maintenance of tubular architecture in the developing testis. Proceedings of the National Academy of Sciences. 2006;103(32):11987–11992. doi:10.1073/pnas.0600994103

65. Tan SWS, Yip GW, Suda T, Baeg GH. Small Maf functions in the maintenance of germline stem cells in the Drosophila testis. Redox Biol. 2018;15:125–134. doi:10.1016/j.redox.2017.12.002

66. Kloth L, Gottlieb A, Helmke B, et al. HMGA2 expression distinguishes between different types of postpubertal testicular germ cell tumour. J Pathol Clin Res. 2015;1(4):239–251. doi:10.1002/cjp2.26

67. Kistler WS, Baas D, Lemeille S, et al. RFX2 Is a Major Transcriptional Regulator of Spermiogenesis. PLoS Genet. 2015;11(7):e1005368. doi:10.1371/journal.pgen.1005368

68. Li L, Dong J, Yan L, et al. Single-Cell RNA-Seq Analysis Maps Development of Human Germline Cells and Gonadal Niche Interactions. Cell Stem Cell. 2017;20(6):858–873.e4. doi:10.1016/j.stem.2017.03.007

69. Rotgers E, Nurmio M, Pietilä E, Cisneros-Montalvo S, Toppari J. E2F1 controls germ cell apoptosis during the first wave of spermatogenesis. Andrology. 2015;3(5):1000–1014. doi:10.1111/andr.12090

70. Maman E, Yung Y, Cohen B, et al. Expression and regulation of sFRP family members in human granulosa cells. Mol Hum Reprod. 2011;17(7):399–404. doi:10.1093/molehr/gar010

71. Der B, Bugacov H, Briantseva BM, McMahon AP. Cadherin adhesion complexes direct cell aggregation in the epithelial transition of Wnt-induced nephron progenitor cells. Development. 2024;151(18):dev202303. doi:10.1242/dev.202303

72. Adam M, Saller S, Ströbl S, et al. Decorin is a part of the ovarian extracellular matrix in primates and may act as a signaling molecule. Hum Reprod. 2012;27(11):3249–3258. doi:10.1093/humrep/des297

73. Myers M, Middlebrook BS, Matzuk MM, Pangas SA. Loss of inhibin alpha uncouples oocyte-granulosa dynamics and disrupts postnatal folliculogenesis. Dev Biol. 2009;334(2):458–467. doi:10.1016/j.ydbio.2009.08.001

74. Okada M, Lee L, Maekawa R, et al. Epigenetic Changes of the Cyp11a1 Promoter Region in Granulosa Cells Undergoing Luteinization During Ovulation in Female Rats. Endocrinology. 2016;157(9):3344–3354. doi:10.1210/en.2016-1264

75. Schaab AM, Stroud KL, Nguyen DT, et al. PAPPA’s role in female reproduction. Reproduction. 2025;170(2):e250012. doi:10.1530/REP-25-0012

76. Cadenas J, Pors SE, Hansen CP, et al. Midkine characterization in human ovaries: potential new variants in follicles. F&S Science. 2023;4(4):294–301. doi:10.1016/j.xfss.2023.09.003

77. Pisarska MD, Barlow G, Kuo FT. Minireview: roles of the forkhead transcription factor FOXL2 in granulosa cell biology and pathology. Endocrinology. 2011;152(4):1199–1208. doi:10.1210/en.2010-1041

78. Liu Q, Kong L, Zhang J, et al. Involvement of GJA1 and Gap Junctional Intercellular Communication between Cumulus Cells and Oocytes from Women with PCOS. Biomed Res Int. 2020;2020:5403904. doi:10.1155/2020/5403904

79. Spitschak M, Hoeflich A. Potential Functions of IGFBP-2 for Ovarian Folliculogenesis and Steroidogenesis. Front Endocrinol. 2018;9. doi:10.3389/fendo.2018.00119

80. Sandell LL, Lynn ML, Inman KE, McDowell W, Trainor PA. RDH10 Oxidation of Vitamin A Is a Critical Control Step in Synthesis of Retinoic Acid during Mouse Embryogenesis. PLOS ONE. 2012;7(2):e30698. doi:10.1371/journal.pone.0030698

81. Workman S, Wilson MJ. RNA sequencing and expression analysis reveal a role for Lhx9 in the haploinsufficient adult mouse ovary. Mol Reprod Dev. 2023;90(5):295–309. doi:10.1002/mrd.23686

82. Dilower I, Niloy AJ, Kumar V, et al. Hedgehog Signaling in Gonadal Development and Function. Cells. 2023;12(3). doi:10.3390/cells12030358

83. Sharma SC, Richards JS. Regulation of AP1 (Jun/Fos) factor expression and activation in ovarian granulosa cells. Relation of JunD and Fra2 to terminal differentiation. J Biol Chem. 2000;275(43):33718–33728. doi:10.1074/jbc.M003555200

84. Southall J, Park S, Choi Y, Jeon H, Ko C, Jo M. Granulosa cell expression of Fos is critical for regulating ovulatory gene expressions in the mouse ovary. FASEB J. 2025;39(4):e70388. doi:10.1096/fj.202402867R

85. Waheed Y, Mojumdar A, Shafiq M, de Marco A, De March M. The fork remodeler helicase-like transcription factor in cancer development: all at once. Biochimica et Biophysica Acta (BBA) - Molecular Basis of Disease. 2024;1870(7):167280. doi:10.1016/j.bbadis.2024.167280

86. Schnabel CA, Selleri L, Cleary ML. Pbx1 is essential for adrenal development and urogenital differentiation. Genesis. 2003;37(3):123–130. doi:10.1002/gene.10235

87. Cadenas J, Pors SE, Hansen CP, et al. Midkine characterization in human ovaries: potential new variants in follicles. F&S Science. 2023;4(4):294–301. doi:10.1016/j.xfss.2023.09.003

88. Elfenbein A, Simons M. Syndecan-4 signaling at a glance. J Cell Sci. 2013;126(17):3799–3804. doi:10.1242/jcs.124636

89. Ogita H, Takai Y. Nectins and nectin-like molecules: Roles in cell adhesion, polarization, movement, and proliferation. IUBMB Life. 2006;58(5-6):334–343. doi:10.1080/15216540600719622

90. Kawasaki A, Torii K, Yamashita Y, et al. Wnt5a promotes adhesion of human dermal fibroblasts by triggering a phosphatidylinositol-3 kinase/Akt signal. Cell Signal. 2007;19(12):2498–2506. doi:10.1016/j.cellsig.2007.07.023

91. Du Y, Han L, Wei H, et al. Granulosa Cell-Secreted KITL Is Involved in Maintaining Zinc Homeostasis in the Oocytes of Neonatal Mouse Ovaries. Antioxidants (Basel). 2025;14(11):1345. doi:10.3390/antiox14111345

92. Hu J, Li S. The Role of Lama2 in Cancer: Current Perspectives. Cancer Res J. 2022;10(4):85–88. doi:10.11648/j.crj.20221004.13

93. Vos MC, van der Wurff AAM, van Kuppevelt TH, Massuger LFAG. The role of MMP-14 in ovarian cancer: a systematic review. J Ovarian Res. 2021;14:101. doi:10.1186/s13048-021-00852-7

94. Vega FM, Ridley AJ. The RhoB small GTPase in physiology and disease. Small GTPases. 2016;9(5):384–393. doi:10.1080/21541248.2016.1253528

95. Yeung CYC, Taylor SH, Garva R, et al. Arhgap28 Is a RhoGAP that Inactivates RhoA and Downregulates Stress Fibers. PLOS ONE. 2014;9(9):e107036. doi:10.1371/journal.pone.0107036

96. Moreno CS. SOX4: The Unappreciated Oncogene. Semin Cancer Biol. 2020;67(Pt 1):57–64. doi:10.1016/j.semcancer.2019.08.027

97. Cole AJ, Panesso-Gómez S, Shah JS, et al. Quiescent ovarian cancer cells secrete follistatin to induce chemotherapy resistance in surrounding cells in response to chemotherapy. Clin Cancer Res. 2023;29(10):1969–1983. doi:10.1158/1078-0432.CCR-22-2254

98. Arimochi J, Ohashi-Kobayashi A, Maeda M. Interaction of Mat-8 (FXYD-3) with Na+/K+-ATPase in Colorectal Cancer Cells. Biological and Pharmaceutical Bulletin. 2007;30(4):648–654. doi:10.1248/bpb.30.648

99. Chen J, Liu J. Characterization of the structure of antithrombin-binding heparan sulfate generated by heparan sulfate 3-*O*-sulfotransferase 5. Biochimica et Biophysica Acta (BBA) - General Subjects. 2005;1725(2):190–200. doi:10.1016/j.bbagen.2005.06.012

100. Farahani RM, Xaymardan M. Platelet-Derived Growth Factor Receptor Alpha as a Marker of Mesenchymal Stem Cells in Development and Stem Cell Biology. Stem Cells Int. 2015;2015:362753. doi:10.1155/2015/362753

101. Richards JS, Ren YA, Candelaria N, Adams JE, Rajkovic A. Ovarian Follicular Theca Cell Recruitment, Differentiation, and Impact on Fertility: 2017 Update. Endocr Rev. 2018;39(1):1–20. doi:10.1210/er.2017-00164

102. Glister C, Satchell L, Bathgate RAD, et al. Functional link between bone morphogenetic proteins and insulin-like peptide 3 signaling in modulating ovarian androgen production. Proceedings of the National Academy of Sciences. 2013;110(15):E1426–E1435. doi:10.1073/pnas.1222216110

103. Hatzirodos N, Hummitzsch K, Irving-Rodgers HF, Rodgers RJ. Transcriptome Profiling of the Theca Interna in Transition from Small to Large Antral Ovarian Follicles. PLOS ONE. 2014;9(5):e97489. doi:10.1371/journal.pone.0097489

104. Kiriakidou M, McAllister JM, Sugawara T, Strauss JF. Expression of steroidogenic acute regulatory protein (StAR) in the human ovary. J Clin Endocrinol Metab. 1996;81(11):4122–4128. doi:10.1210/jcem.81.11.8923870

105. Lardenois A, Suglia A, Moore C, et al. Single-cell transcriptome landscape of developing fetal gonads defines somatic cell lineage specification in humans. bioRxiv. Preprint posted online August 8, 2023:2023.08.07.552336. doi:10.1101/2023.08.07.552336

106. Kakuta H, Iguchi T, Sato T. The Involvement of Granulosa Cells in the Regulation by Gonadotropins of Cyp17a1 in Theca Cells. In Vivo. 2018;32(6):1387–1401. doi:10.21873/invivo.11391

107. Liu C, Peng J, Matzuk MM, Yao HHC. Lineage Specification of Ovarian Theca Cells Requires Multi-Cellular Interactions via Oocyte and Granulosa Cells. Nat Commun. 2015;6:6934. doi:10.1038/ncomms7934

108. Choi Y, Jeon H, Akin JW, Curry TE, Jo M. The FOS/AP-1 Regulates Metabolic Changes and Cholesterol Synthesis in Human Periovulatory Granulosa Cells. Endocrinology. 2021;162(9):bqab127. doi:10.1210/endocr/bqab127

109. Tan HJ, Deng ZH, Shen H, Deng HW, Xiao HM. Single-cell RNA-seq identified novel genes involved in primordial follicle formation. Front Endocrinol (Lausanne). 2023;14:1285667. doi:10.3389/fendo.2023.1285667

110. Samanta D, Almo SC. Nectin family of cell-adhesion molecules: structural and molecular aspects of function and specificity. Cell Mol Life Sci. 2014;72(4):645–658. doi:10.1007/s00018-014-1763-4

111. Kawagishi R, Tahara M, Morishige K, et al. Expression of nectin-2 in mouse granulosa cells. Eur J Obstet Gynecol Reprod Biol. 2005;121(1):71–76. doi:10.1016/j.ejogrb.2004.12.019

112. Hansen CL, Pelegri F. Primordial Germ Cell Specification in Vertebrate Embryos: Phylogenetic Distribution and Conserved Molecular Features of Preformation and Induction. Front Cell Dev Biol. 2021;9:730332. doi:10.3389/fcell.2021.730332

113. Bächler M, Menshykau D, De Geyter Ch, Iber D. Species-specific differences in follicular antral sizes result from diffusion-based limitations on the thickness of the granulosa cell layer. Mol Hum Reprod. 2014;20(3):208–221. doi:10.1093/molehr/gat078

114. Ridley A. Rho Gtpases. J Cell Biol. 2000;150(4):f107-f109. doi:10.1083/jcb.150.4.f107

115. Abarca-Buis RF, Mandujano-Tinoco EA, Krötzsch E. The complexity of TGFβ/activin signaling in regeneration. J Cell Commun Signal. 2021;15(1):7–23. doi:10.1007/s12079-021-00605-7

116. Abdelaal T, Michielsen L, Cats D, et al. A comparison of automatic cell identification methods for single-cell RNA sequencing data. Genome Biol. 2019;20:194. doi:10.1186/s13059-019-1795-z

117. Lyons A, Brown J, Davenport KM. Single-Cell Sequencing Technology in Ruminant Livestock: Challenges and Opportunities. Curr Issues Mol Biol. 2024;46(6):5291–5306. doi:10.3390/cimb46060316

