## Supplementary material for "Comparative single-cell transcriptomic roadmap of mammalian fetal ovarian development": suppl. figures

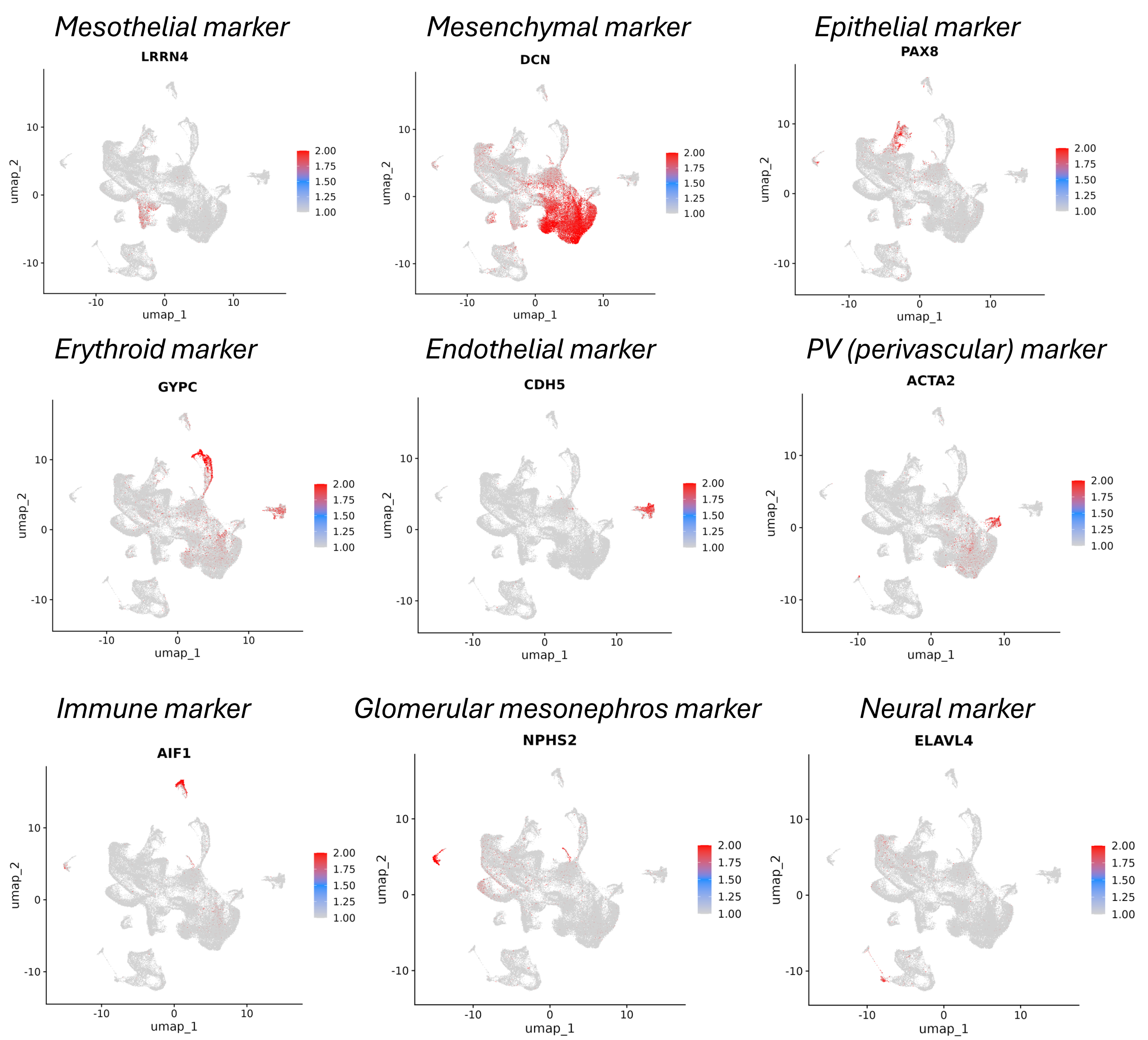


**Supplementary Figure 1. Validation of cell type annotations using canonical marker genes**

Feature plots showing the expression of representative marker genes used to identify major ovarian cell populations in the integrated single-cell dataset. Markers include *LRRN4* for mesothelial cells, *DCN* for mesenchymal cells, *PAX8* for epithelial cells, *GYPC* for erythroid cells, *CDH5* for endothelial cells, *ACTA2* for perivascular (PV) cells, *AIF1* for immune cells, *NPHS2* for glomerular mesonephros cells, and *ELAVL4* for neural cells. Gene expression levels are overlaid on the UMAP embedding.

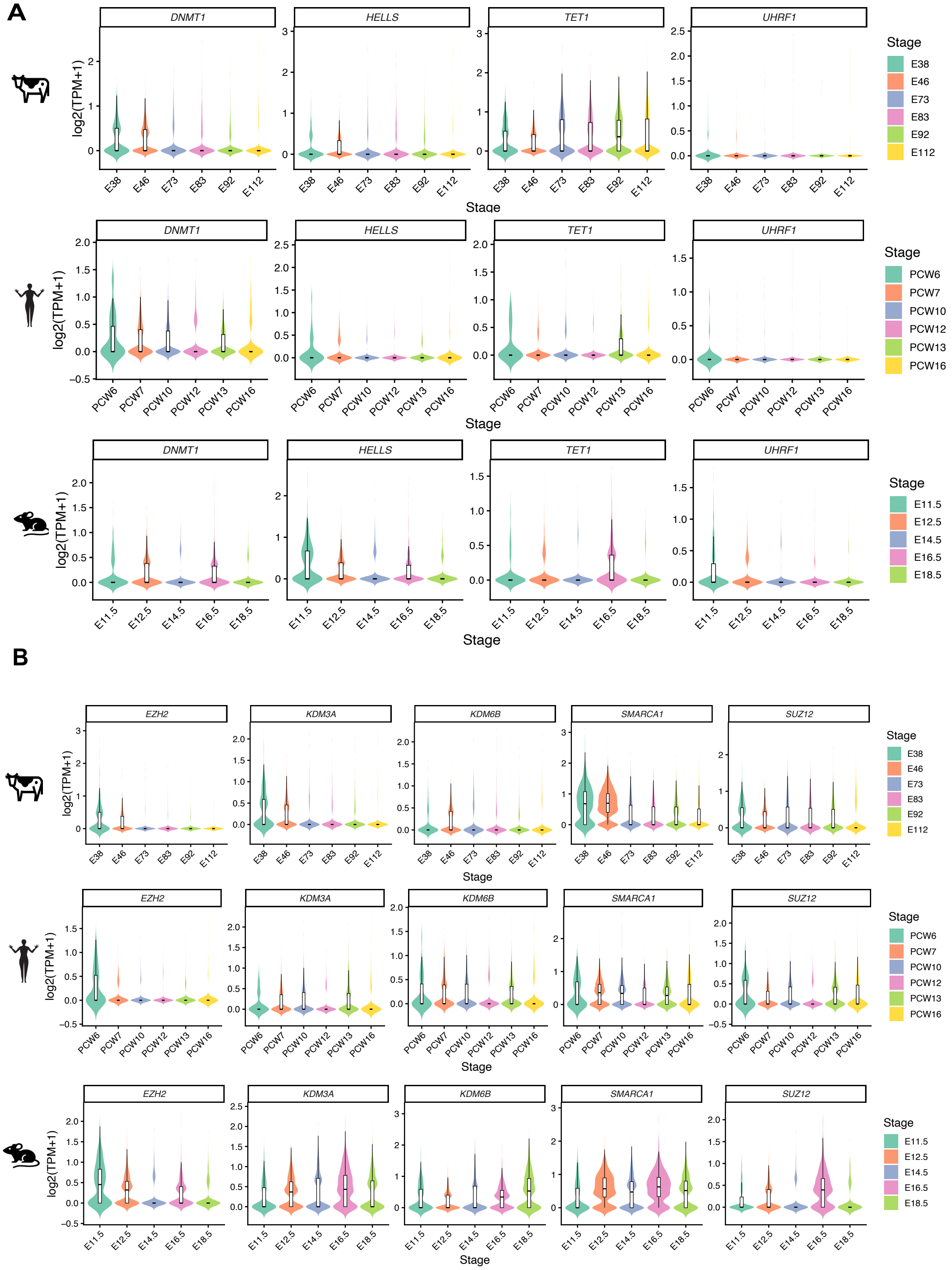
A
DNMT1
HELLS
TET1
UHRF1
log2(TPM+1)
Stage
E38
E46
E73
E83
E92
E112
PCW6
PCW7
PCW10
PCW12
PCW13
PCW16
E11.5
E12.5
E14.5
E16.5
E18.5
B
EZH2
KDM3A
KDM6B
SMARCA1
SUZ12

### Supplementary Figure 2. Cross-species dynamics of epigenetic regulators during early granulosa cell development

(A) Expression dynamics of DNA methylation–related regulators (*DNMT1*, *HELLS*, *TET1*, *UHRF1*) across developmental stages in bovine (E38-E112), human (PCW6–PCW16) and mouse (E11.5–E18.5) granulosa cells. (B) Expression dynamics of chromatin remodeling and histone modification–related regulators (*EZH2*, *KDM3A*, *KDM6B*, *SMARCA1*, *SUZ12*) across developmental stages in bovine (E38–E112), human (PCW6–PCW16), and mouse (E11.5–E18.5) granulosa cells. Violin plots show the distribution of log2(TPM + 1) expression levels for each gene at each developmental stage, with embedded boxplots indicating median and interquartile range. Colors represent developmental stages as indicated. Species are denoted by icons on the left (cattle, human, and mouse).

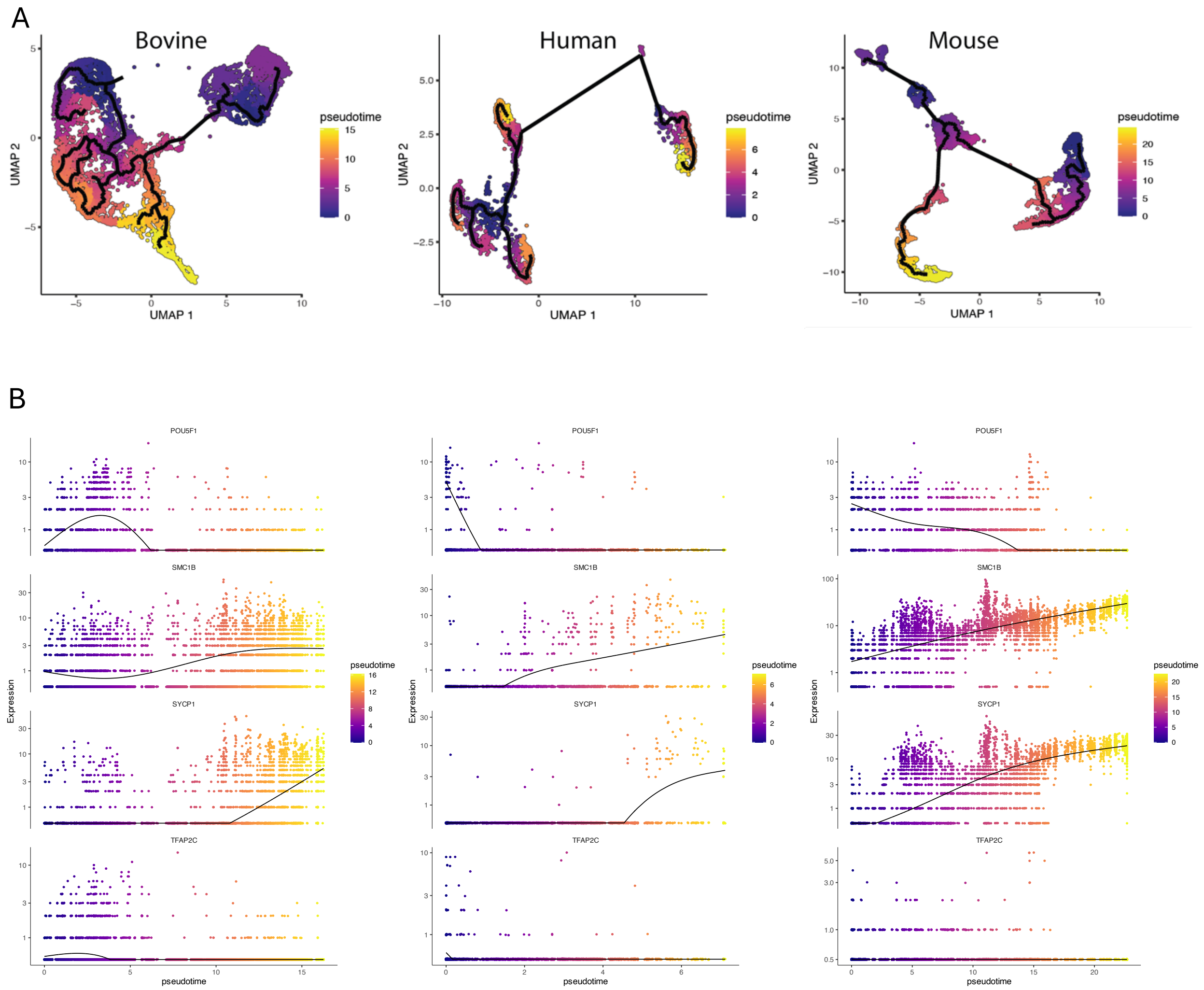


**Supplementary Figure 3. Pseudotime trajectory and dynamic gene expression during female germ cell development across species**

Pseudotime trajectory analysis of germ cells from bovine, human, and mouse. (A) UMAP embeddings overlaid with inferred developmental trajectories, with cells colored by pseudotime from early to late stages. (B) Expression dynamics of representative genes plotted along pseudotime in each species. Gene expression levels are shown for individual cells, with smoothed curves indicating overall trends. Selected genes illustrate stage-dependent transcriptional changes during female germ cell progression.

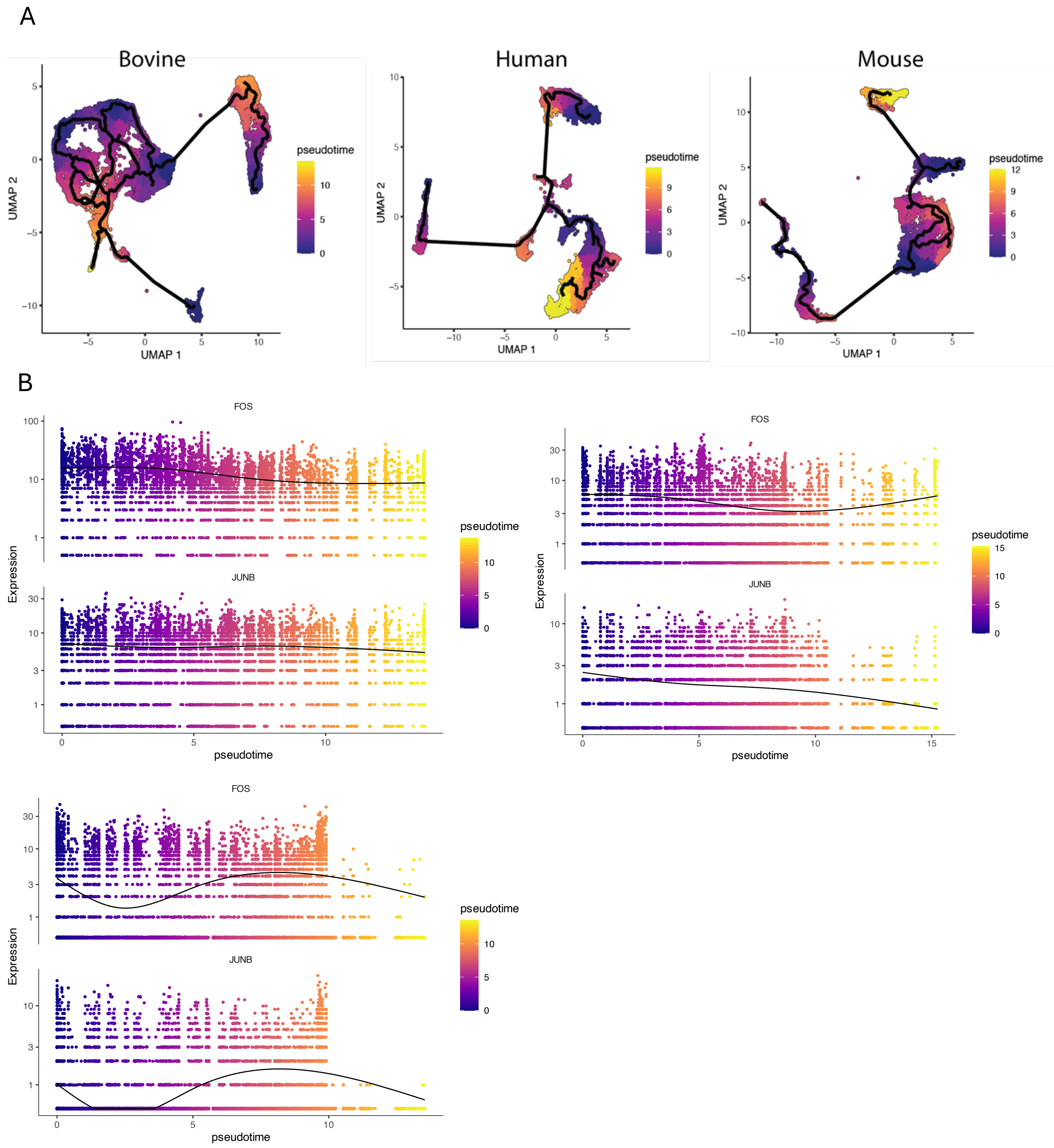


**Supplementary Figure 4. Pseudotime trajectory and dynamic gene expression during granulosa cell development across species**

Pseudotime trajectory analysis of granulosa cells from bovine, human, and mouse. (A) UMAP embeddings overlaid with inferred developmental trajectories, with cells colored by pseudotime from early to late stages. (B) Expression dynamics of representative genes plotted along pseudotime in each species. Each point represents an individual cell, and smoothed curves indicate overall expression trends, illustrating stage-dependent transcriptional changes during granulosa cell development.

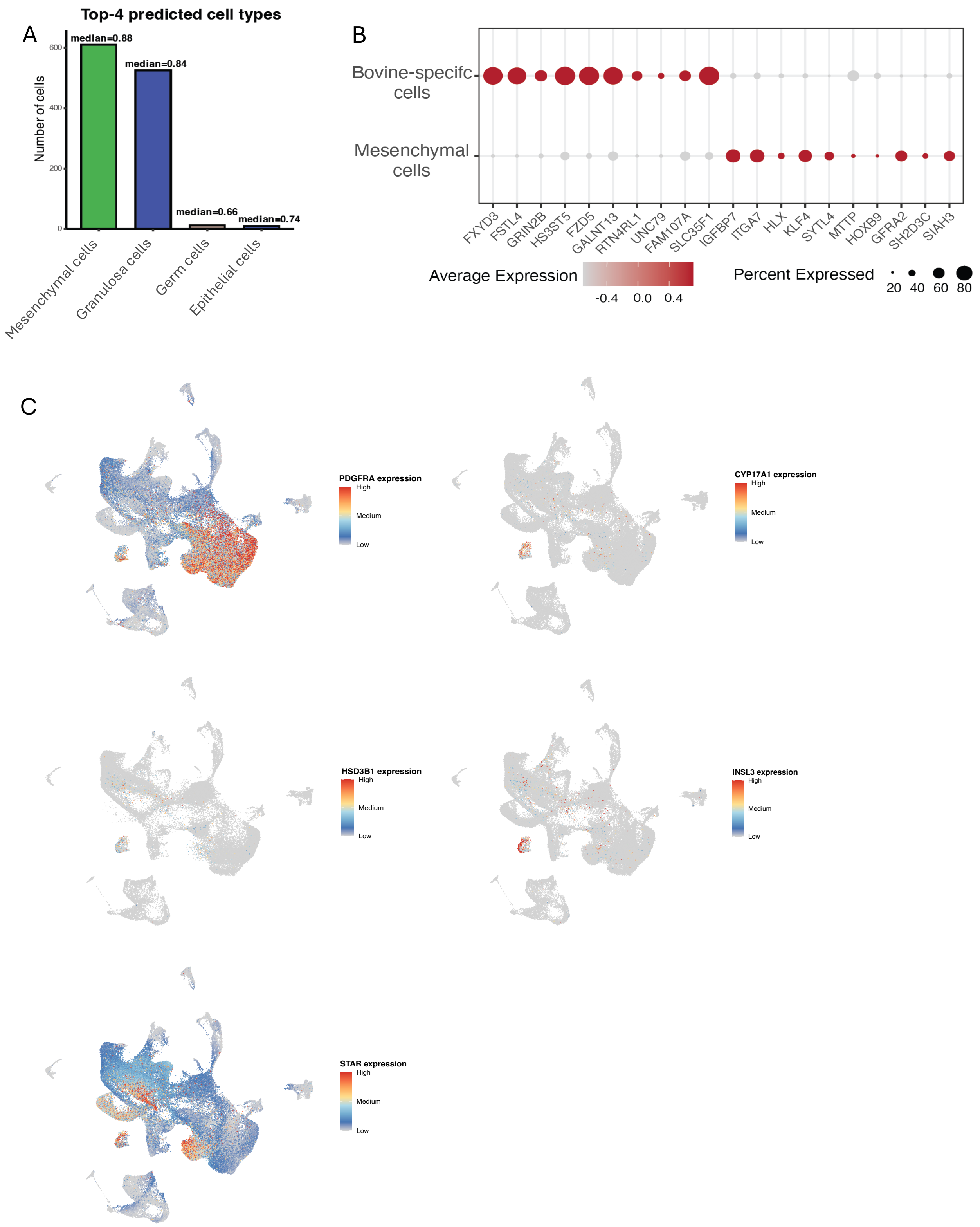
A
Top-4 predicted cell types
Number of cells
600
400
200
0
median=0.88
median=0.84
median=0.66
median=0.74
Mesenchymal cells
Granulosa cells
Germ cells
Epithelial cells
B
Bovine-specifc cells
Mesenchymal cells
FXYD3
FSTL4
GRIN2B
HS3ST5
FZD5
GALNT13
RTN4RL1
UNC79
FAM107A
SLC35F1
IGFBP7
ITGA7
HLX
KLF4
SYTL4
MTTP
HOXB9
GFRA2
SH2D3C
SIAH3
Average Expression
-0.4
0.0
0.4
Percent Expressed
20
40
60
80
C
PDGFRA expression
CYP17A1 expression
HSD3B1 expression
INSL3 expression
STAR expression
High
Medium
Low

**Supplementary Figure 5. Characterization of a bovine-specific somatic cell population using SVM classification and marker expression**

(A) Prediction of the cell type identity of the bovine-specific cell population using an SVM classifier trained on all other bovine ovarian cell types. Bar plots show the number of bovine-specific cells assigned to the top four predicted cell types, with median prediction probabilities indicated. Mesenchymal and granulosa cells accounted for the majority of predictions, whereas smaller fractions were classified as germ cells or epithelial cells. (B) Dot plot showing differential gene expression between the bovine-specific cell population and mesenchymal cells. Dot color indicates average scaled expression, and dot size represents the percentage of cells expressing each gene. (C) Feature plots showing the expression of the representative mesenchymal marker *PDGFRA* and theca cell–associated markers (*CYP17A1*, *HSD3B1*, *INSL3*, and *STAR*) projected onto the UMAP embedding. The bovine-specific cell population shows partial expression of steroidogenic markers.
